# Variant Impact Predictor database (VIPdb), version 2: Trends from 25 years of genetic variant impact predictors

**DOI:** 10.1101/2024.06.25.600283

**Authors:** Yu-Jen Lin, Arul S. Menon, Zhiqiang Hu, Steven E. Brenner

**Author notes:** Yu-Jen Lin and Arul S. Menon are joint first authors. Steven E. Brenner is the corresponding author. **Correspondence**: Steven E. Brenner Department of Plant and Microbial Biology, 111 Koshland Hall #3102 University of California Berkeley, CA 94720-3102, USA. **Email Yu-Jen Lin:**, **Arul S. Menon:**, **Zhiqiang Hu:**, **Steven E. Brenner:**.

## Abstract

**Background:** Variant interpretation is essential for identifying patients’ disease-causing genetic variants amongst the millions detected in their genomes. Hundreds of Variant Impact Predictors (VIPs), also known as Variant Effect Predictors (VEPs), have been developed for this purpose, with a variety of methodologies and goals. To facilitate the exploration of available VIP options, we have created the Variant Impact Predictor database (VIPdb).

**Results:** The Variant Impact Predictor database (VIPdb) version 2 presents a collection of VIPs developed over the past 25 years, summarizing their characteristics, ClinGen calibrated scores, CAGI assessment results, publication details, access information, and citation patterns. We previously summarized 217 VIPs and their features in VIPdb in 2019. Building upon this foundation, we identified and categorized an additional 186 VIPs, resulting in a total of 403 VIPs in VIPdb version 2. The majority of the VIPs have the capacity to predict the impacts of single nucleotide variants and nonsynonymous variants. More VIPs tailored to predict the impacts of insertions and deletions have been developed since the 2010s. In contrast, relatively few VIPs are dedicated to the prediction of splicing, structural, synonymous, and regulatory variants. The increasing rate of citations to VIPs reflects the ongoing growth in their use, and the evolving trends in citations reveal development in the field and individual methods.

**Conclusions:** VIPdb version 2 summarizes 403 VIPs and their features, potentially facilitating VIP exploration for various variant interpretation applications.

**Availability:** VIPdb version 2 is available at https://genomeinterpretation.org/vipdb

## Background

Advances in sequencing technologies, including gene panels, whole exome sequencing, whole genome sequencing, and long read sequencing, have revolutionized the investigation of genetic variation on a large scale and hence have accelerated the discovery of novel genetic etiologies of diseases and improved the efficiency of diagnosis (1, 2). Typically, thousands to millions of variants are identified in each individual (3, 4), making it challenging to distinguish disease-causing variants from non-contributory ones. Consequently, methods to predict the impacts of variants being disease-causing are essential (5, 6).

This need prompted the development of Variant Impact Predictors (VIPs), tools or databases designed to predict the consequences of genetic variants. Hundreds of genetic VIPs have been developed, with a variety of methodologies and goals (7). Some overlapping categories of variants considered by different tools are single nucleotide variations (SNVs), insertions and deletions (indels), structural variations (SVs), nonsynonymous variants, synonymous variants, splicing variants, and regulatory variants. VIPs are designed for different contexts, such as for germline variants, somatic variants, or specific diseases or genes. The variety of VIPs underscores the complex nature of variant interpretation and poses a challenge for users in identifying the most suitable VIPs for their specific needs.

Many computational impact prediction methods have been developed, yet the field lacks a clear consensus on their appropriate use and interpretation (8). Recognizing the need for an organized approach to explore available VIPs, several research entities have constructed resources facilitating the informed use of VIPs. Initiatives like the Critical Assessment of Genome Interpretation (CAGI) conduct community experiments to assess VIPs across different variant types and contexts (8, 9, 10). The dbNSFP (database for Nonsynonymous Single-nucleotide polymorphisms’ Functional Predictions) hosts precomputes of several VIP results (11). OpenCRAVAT integrates hundreds of VIP analyses of cancer-related variants in one platform, enhancing accessibility for users (12). These resources have played an important role in introducing users to VIP options. Consequently, we developed VIPdb to serve as a comprehensive resource for exploring VIPs.

To systematically evaluate the pathogenicity of a variant in a clinical laboratory, ACMG/AMP has established guidelines for interpreting genetic variants that integrate several lines of evidence, including population data, functional data, segregation data, and computational prediction (13). Historically, VIPs provided only supporting evidence in determining the pathogenicity or benignity of variants in clinical settings. However, recent ClinGen clinical recommendations allow VIPs the potential to provide stronger evidence (14). This greater role for VIPs in providing evidence for clinical decisions could improve genetic disease diagnosis.

The Variant Impact Predictor database (VIPdb) offers a curation of available computational tools for predicting variant impact. Initially established in 2007 and 2010 (15), the database was last updated in 2019 (7). VIPdb version 2 is a comprehensive update through January 2, 2024.

## Implementation

Our identification of VIPs involved searching for potential VIPs and examining their articles to determine whether they should be included in VIPdb. In the initial step, we searched the literature using the query “(((tool(Title]) OR (pipeline(Title])) AND (variant(Title/Abstract]))” on PubMed and collected potential VIPs citing pioneering VIPs (SIFT, PolyPhen, ANNOVAR, and SnpEff) (16, 17, 18, 19, 20, 21, 22, 23, 24, 25, 26, 27). Additionally, we gathered potential VIPs from existing databases such as OpenCRAVAT and dbNSFP, as well as from submissions by VIP developers. Subsequently, we examined the literature and included only programs capable of handling variant data, such as VCF files, rsID, or location in the genome, and providing evidence or predictions of the variant impacts. Overall, this resulted in the identification of 186 additional VIPs, augmenting the VIPdb to a total of 403 VIPs (16, 17, 19, 21, 22, 23, 24, 25, 26, 27, 28, 29, 30, 31, 32, 33, 34, 35, 36, 37, 38, 39, 40, 41, 42, 43, 44, 45, 46, 47, 48, 49, 50, 51, 52, 53, 54, 55, 56, 57, 58, 59, 60, 61, 62, 63, 64, 65, 66, 67, 68, 69, 70, 71, 72, 73, 74, 75, 76, 77, 78, 79, 80, 81, 82, 83, 84, 85, 86, 87, 88, 89, 90, 91, 92, 93, 94, 95, 96, 97, 98, 99, 100, 101, 102, 103, 104, 105, 106, 107, 108, 109, 110, 111, 112, 113, 114, 115, 116, 117, 118, 119, 120, 121, 122, 123, 124, 125, 126, 127, 128, 129, 130, 131, 132, 133, 134, 135, 136, 137, 138, 139, 140, 141, 142, 143, 144, 145, 146, 147, 148, 149, 150, 151, 152, 153, 154, 155, 156, 157, 158, 159, 160, 161, 162, 163, 164, 165, 166, 167, 168, 169, 170, 171, 172, 173, 174, 175, 176, 177, 178, 179, 180, 181, 182, 183, 184, 185, 186, 187, 188, 189, 190, 191, 192, 193, 194, 195, 196, 197, 198, 199, 200, 201, 202, 203, 204, 205, 206, 207, 208, 209, 210, 211, 212, 213, 214, 215, 216, 217, 218, 219, 220, 221, 222, 223, 224, 225, 226, 227, 228, 229, 230, 231, 232, 233) (11, 18, 20, 234, 235, 236, 237, 238, 239, 240, 241, 242, 243, 244, 245, 246, 247, 248, 249, 250, 251, 252, 253, 254, 255, 256, 257, 258, 259, 260, 261, 262, 263, 264, 265, 266, 267, 268, 269, 270, 271, 272, 273, 274, 275, 276, 277, 278, 279, 280, 281, 282, 283, 284, 285, 286, 287, 288, 289, 290, 291, 292, 293, 294, 295, 296, 297, 298, 299, 300, 301, 302, 303, 304, 305, 306, 307, 308, 309, 310, 311, 312, 313, 314, 315, 316, 317, 318, 319, 320, 321, 322, 323, 324, 325, 326, 327, 328, 329, 330, 331, 332, 333, 334, 335, 336, 337, 338, 339, 340, 341, 342, 343, 344, 345, 346, 347, 348, 349, 350, 351, 352, 353, 354, 355, 356, 357, 358, 359, 360, 361, 362, 363, 364, 365, 366, 367, 368, 369, 370, 371, 372, 373, 374, 375, 376, 377, 378, 379, 380, 381, 382, 383, 384, 385, 386, 387, 388, 389, 390, 391, 392, 393, 394, 395, 396, 397, 398, 399, 400, 401, 402, 403, 404, 405, 406, 407, 408, 409, 410, 411, 412, 413, 414, 415).

To facilitate users’ exploration of available VIPs, we described key features of each VIP. VIPs primarily designed for variant impact prediction were labeled as such. VIPs not originally designed for variant impact prediction but nonetheless used for this purpose, such as those estimating conservation scores and population allele frequencies, were categorized as non-primary. VIPs containing clinical classifications, functional data, or population data were categorized as databases, whereas VIPs utilizing databases for computing variant impact predictions were classified as non-databases. Furthermore, as VIPs are designed for different types of genetic variants, we classified the VIPs according to the following overlapping categories of input: single nucleotide variant (SNV), insertion and deletion (indel) variant, structural variant (SV), nonsynonymous/nonsense variant, synonymous variant, splicing variant, and regulatory region variants, with some overlap among these categories. Licensing information, including whether the VIP is free for academic or commercial use, was also included. In addition, we provided details about accessing VIPs, such as homepage links and source code availability.

In VIPdb version 2, we have made enhancements to inform clinical decision-making. We incorporated calibrated threshold scores recommended by ClinGen for clinical use (14) with ACMG/AMP guidelines for variant classification (13). Additionally, we included community assessment results from the CAGI 6 Annotate All Missense / Missense Marathon challenge (416) to enable users to compare the overall performance of methods and the performance on subsets with high specificity or high sensitivity.

To understand the trends of genetic VIPs over 25 years, we conducted a citation analysis. We utilized the Entrez module in Biopython to retrieve citation information from the PubMed database. Specifically, the elink function was employed to collect the number of articles citing each VIP, and the esummary function allowed for the collection of publication years for these citations. These functions facilitated the automatic collection of citation numbers by year for each VIP.

In summary, VIPdb version 2 presents a collection of 403 VIPs developed over the last 25 years, with their characteristics, citation patterns, publication details, and access information. VIPdb version 2 is publicly accessible at https://genomeinterpretation.org/vipdb

## Results

We incorporated 186 additional VIPs into VIPdb version 2, alongside the existing 217 VIPs in the previous version of VIPdb. We summarized the characteristics of the 403 VIPs in VIPdb version 2. Among the 403 VIPs in VIPdb version 2, 274 are core VIPs, defined as VIPs primarily designed for variant impact prediction and not a database.

An analysis of the variant type used by VIP showed a predominant focus on predicting the impacts of single nucleotide variants (SNVs) and nonsynonymous variants (Fig. 1). Since the 2010s, there has been a notable surge in the development of VIPs tailored for insertions and deletions (indels), while VIPs dedicated to predicting the impacts of splicing, structural, synonymous, and regulatory variants have grown more modestly (Fig. 1). These observations about VIP variant type not only highlight current focus on but also identify areas that have been less explored, suggesting potential directions for future research.

**Figure. 1.**
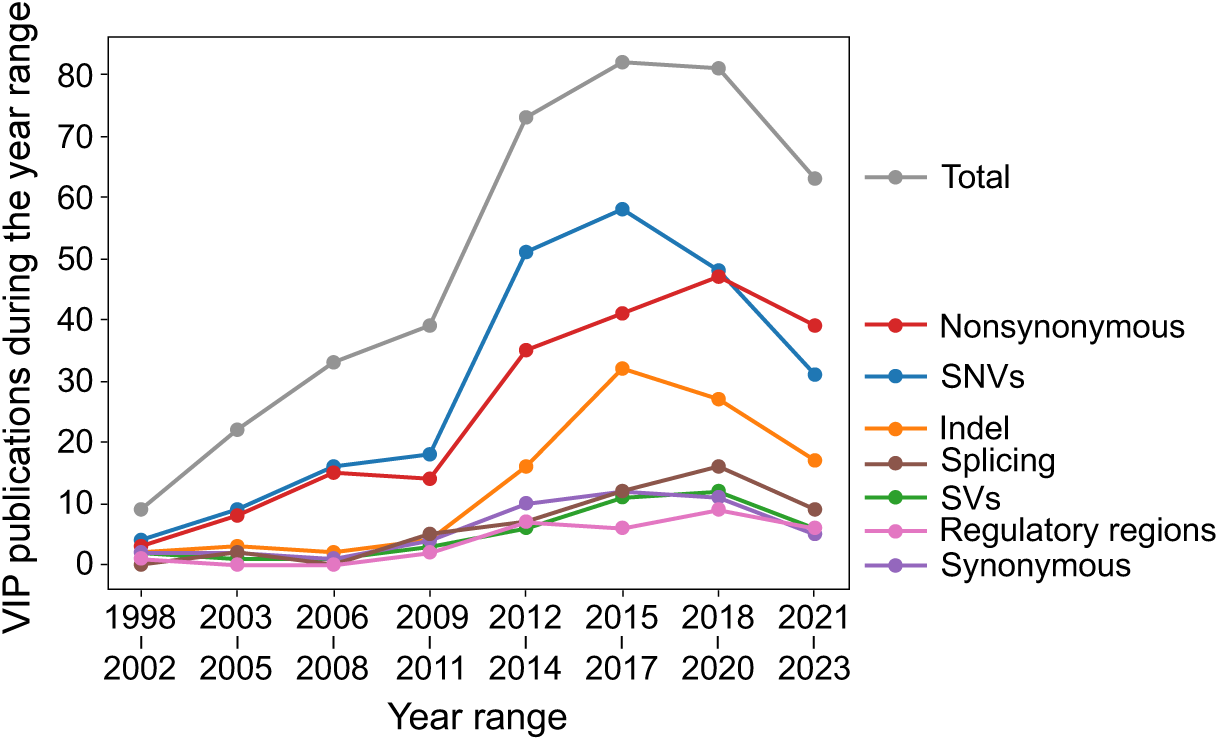
VIP variant type focus.

The citation rate of VIPs continues to rise, while the annual publications of VIPs have reached a plateau (Fig. 2). The increasing citation rates for both the 274 core VIPs and the 129 non-core VIPs reflect the ongoing growth of VIP usage (Fig. 2A). The median total citation for VIPs is 41 from 1998 to 2023, with a 95% quantile of 2612 citations (Fig. 2B). Annual publication showed a stabilization in VIP publications, with some being subsequent publications from previous work (Fig. 2C).

**Figure 2.**
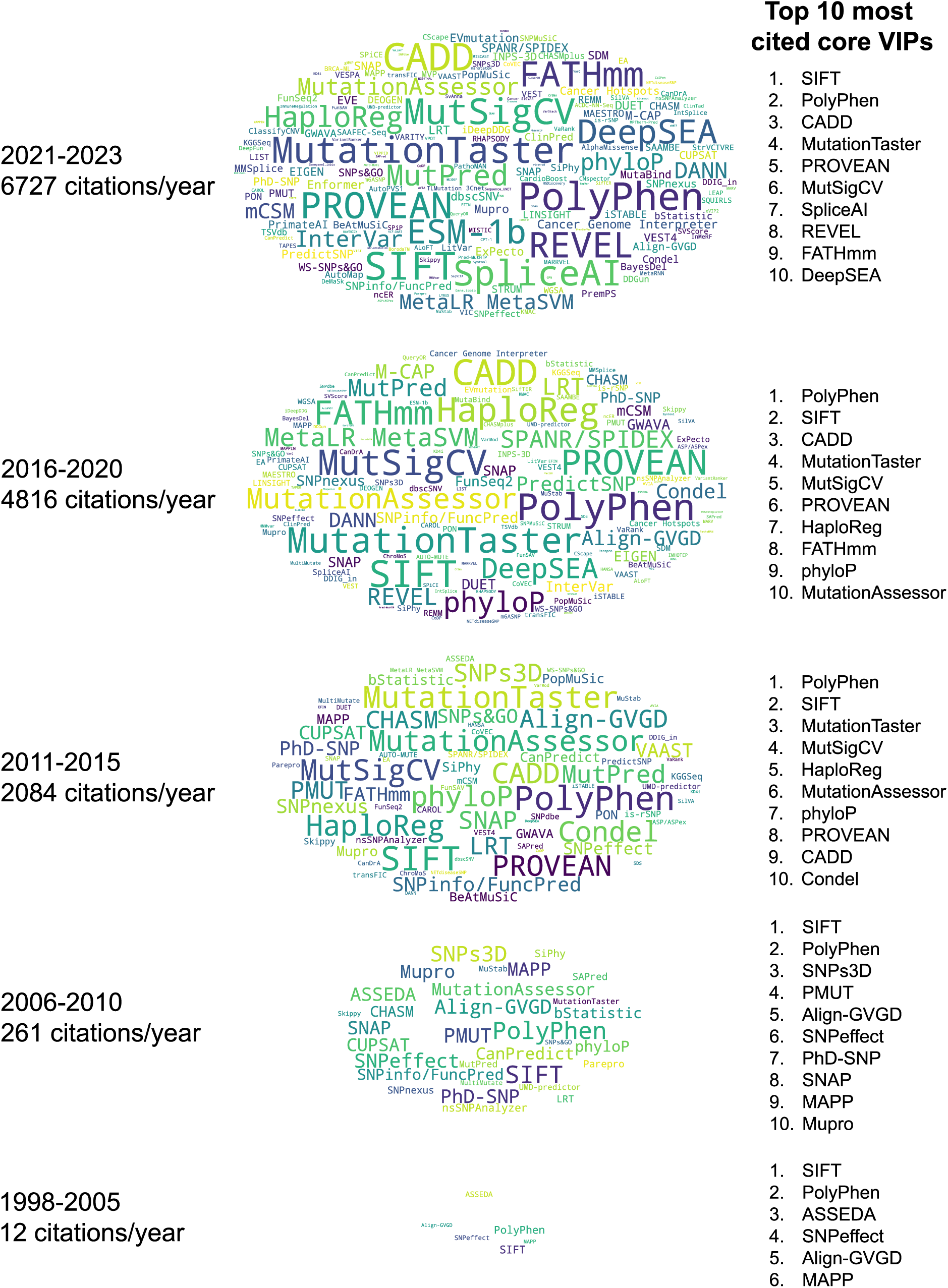
Citation and publication analysis of 403 VIPs. (a) Citations each year for 274 core VIPs (blue) and 126 non-core VIPs (gray). (b) Histogram of total citations for core VIPs (blue) and non-core VIPs (gray). (c) VIPs published per year, with original publications in light blue (core) and light gray (non-core), and subsequent publications in dark blue (core) and dark gray (non-core).

The citation trend of 274 core VIPs from 1998 to 2023 is shown in Fig. 3 and 4. The citation analysis revealed that SIFT and PolyPhen, among the earliest, are the most cited core VIPs (Fig. 3 and 4).

**Figure 3.**
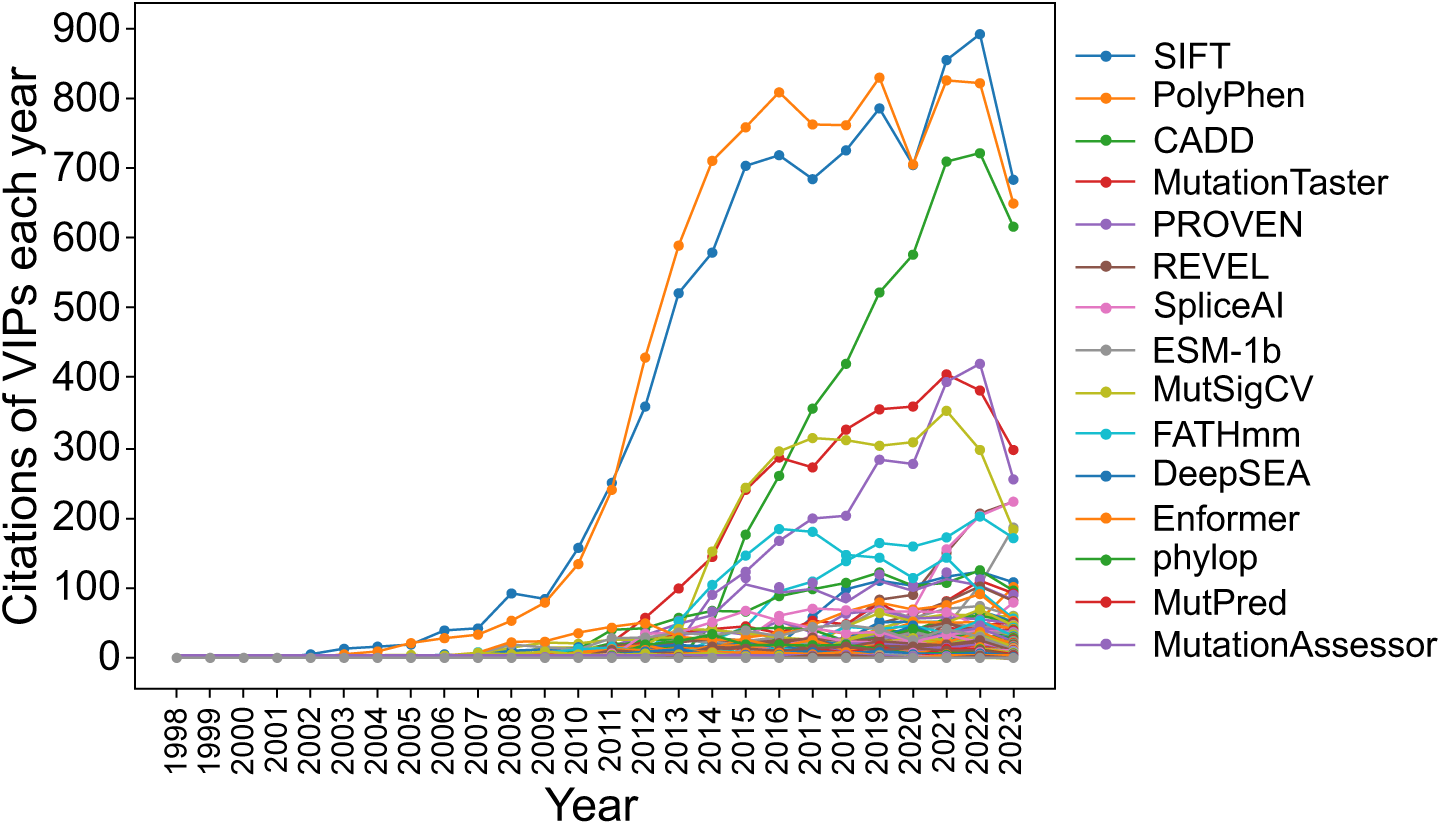
Citation trend of 274 core VIPs (1998 to 2023). Word clouds representing core VIPs over a specific time period, using cumulative citations for core VIPs with multiple publications. Font sizes in the word clouds correspond to the logarithm of citation counts for each period, and cloud heights are scaled by the logarithm of the annual citation averages. The top 10 most cited core VIPs during the period are listed. Note: Core VIPs are methods primarily designed for variant impact prediction and are not classified as databases.

**Figure 4.**
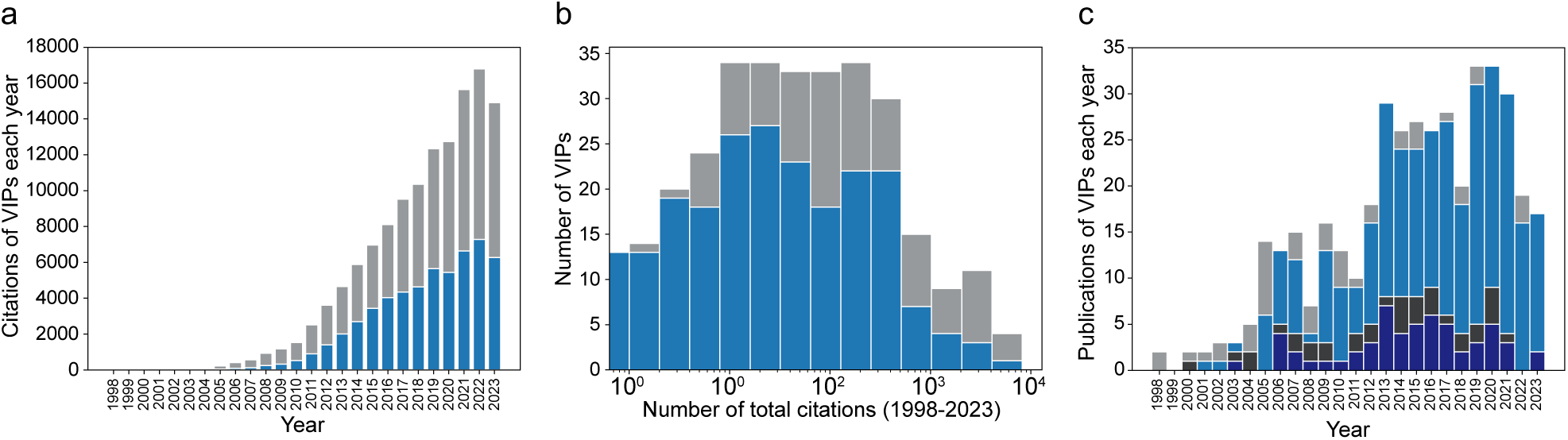
Citation trend of the top 15 most cited core VIPs in the year 2023. Note: Core VIPs are methods primarily designed for variant impact prediction and are not classified as databases.

## Discussion and Conclusions

VIPdb version 2 provides a comprehensive view of VIPs. To identify the most appropriate VIPs for user’s specific needs, users are advised to thoroughly assess the strengths and weaknesses of VIPs before determining their suitability for use. For example, initiatives like the Critical Assessment of Genome Interpretation (CAGI) conduct community experiments to assess VIPs across different variant types and contexts (8, 9, 10).

With 403 curated VIPs, VIPdb version 2 provides a comprehensive overview of programs designed for variant impact prediction, along with their characteristics, citation patterns, publication details, and access information. VIPdb version 2 is available on the CAGI website (https://genomeinterpretation.org/vipdb). We invite submissions of new VIPs to the next version of VIPdb.

## Availability and requirements

Project name: Variant Impact Predictor Database (VIPdb) Project home page: https://genomeinterpretation.org/vipdb

Operating system(s): Platform independent Programming language: Not applicable Other requirements: Not applicable

Any restrictions to use by non-academics: Not applicable

VIPdb version 2 is available at https://genomeinterpretation.org/vipdb

## List of abbreviations

VIP: Variant Impact Predictor
VIPdb: Variant Impact Predictor Database

## Declarations

Ethics approval and consent to participate: Not applicable

Consent for publication: Not applicable

## Availability of data and materials

The VIPdb database is available on the CAGI website (https://genomeinterpretation.org/vipdb), and the full version can be downloaded as a spreadsheet.

## Authors’ contributions

YL conducted summary and visualization analyses and wrote the manuscript. AS, under the guidance of YL, curated VIPs and drafted the manuscript. ZH provided initial advisory input to YL and AS. SEB oversaw the project, provided supervision to YL, AS, and ZH, and edited the manuscript.

## Acknowledgments

We gratefully acknowledge Changhua Yu, Mabel Furutsuki, Gaia Andreoletti, Melissa Ly, Roger Hoskins, and Aashish N Adhikari for their work on VIPdb version 1 (7). We thank Changhua Yu also for his initial guidance in curating VIPdb.

## References

1. Marwaha S, Knowles JW, Ashley EA. A guide for the diagnosis of rare and undiagnosed disease: beyond the exome. Genome Med. 2022;14(1):23.

2. Schobers G, Derks R, den Ouden A, Swinkels H, van Reeuwijk J, Bosgoed E, et al. Genome sequencing as a generic diagnostic strategy for rare disease. Genome Med. 2024;16(1):32.

3. Fowler DM, Adams DJ, Gloyn AL, Hahn WC, Marks DS, Muffley LA, et al. An Atlas of Variant Effects to understand the genome at nucleotide resolution. Genome Biol. 2023;24(1):147.

4. Marian AJ. Clinical Interpretation and Management of Genetic Variants. JACC Basic Transl Sci. 2020;5(10):1029–42.

5. Papadimitriou S, Gazzo A, Versbraegen N, Nachtegael C, Aerts J, Moreau Y, et al. Predicting disease-causing variant combinations. Proc Natl Acad Sci U S A. 2019;116(24):11878–87.

6. Wang D, Li J, Wang Y, Wang E. A comparison on predicting functional impact of genomic variants. NAR Genom Bioinform. 2022;4(1):lqab122.

7. Hu Z, Yu C, Furutsuki M, Andreoletti G, Ly M, Hoskins R, et al. VIPdb, a genetic Variant Impact Predictor Database. Hum Mutat. 2019;40(9):1202–14.

8. Andreoletti G, Pal LR, Moult J, Brenner SE. Reports from the fifth edition of CAGI: The Critical Assessment of Genome Interpretation. Hum Mutat. 2019;40(9):1197–201.

9. Critical Assessment of Genome Interpretation C. CAGI, the Critical Assessment of Genome Interpretation, establishes progress and prospects for computational genetic variant interpretation methods. Genome Biol. 2024;25(1):53.

10. Hoskins RA, Repo S, Barsky D, Andreoletti G, Moult J, Brenner SE. Reports from CAGI: The Critical Assessment of Genome Interpretation. Hum Mutat. 2017;38(9):1039–41.

11. Liu X, Li C, Mou C, Dong Y, Tu Y. dbNSFP v4: a comprehensive database of transcript-specific functional predictions and annotations for human nonsynonymous and splice-site SNVs. Genome Med. 2020;12(1):103.

12. Pagel KA, Kim R, Moad K, Busby B, Zheng L, Tokheim C, et al. Integrated Informatics Analysis of Cancer-Related Variants. JCO Clin Cancer Inform. 2020;4:310–7.

13. Richards S, Aziz N, Bale S, Bick D, Das S, Gastier-Foster J, et al. Standards and guidelines for the interpretation of sequence variants: a joint consensus recommendation of the American College of Medical Genetics and Genomics and the Association for Molecular Pathology. Genet Med. 2015;17(5):405–24.

14. Pejaver V, Byrne AB, Feng BJ, Pagel KA, Mooney SD, Karchin R, et al. Calibration of computational tools for missense variant pathogenicity classification and ClinGen recommendations for PP3/BP4 criteria. Am J Hum Genet. 2022;109(12):2163–77.

15. Brenner SE. Common sense for our genomes. Nature. 2007;449(7164):783-4.

16. Hu J, Ng PC. SIFT Indel: predictions for the functional effects of amino acid insertions/deletions in proteins. PLoS One. 2013;8(10):e77940.

17. Kumar P, Henikoff S, Ng PC. Predicting the effects of coding non-synonymous variants on protein function using the SIFT algorithm. Nat Protoc. 2009;4(7):1073–81.

18. Ng PC, Henikoff S. SIFT: Predicting amino acid changes that affect protein function. Nucleic Acids Res. 2003;31(13):3812–4.

19. Sim NL, Kumar P, Hu J, Henikoff S, Schneider G, Ng PC. SIFT web server: predicting effects of amino acid substitutions on proteins. Nucleic Acids Res. 2012;40(Web Server issue):W452-7.

20. Vaser R, Adusumalli S, Leng SN, Sikic M, Ng PC. SIFT missense predictions for genomes. Nat Protoc. 2016;11(1):1–9.

21. Adzhubei I, Jordan DM, Sunyaev SR. Predicting functional effect of human missense mutations using PolyPhen-2. Curr Protoc Hum Genet. 2013;Chapter 7:Unit7 20.

22. Adzhubei IA, Schmidt S, Peshkin L, Ramensky VE, Gerasimova A, Bork P, et al. A method and server for predicting damaging missense mutations. Nat Methods. 2010;7(4):248–9.

23. Ramensky V, Bork P, Sunyaev S. Human non-synonymous SNPs: server and survey. Nucleic Acids Res. 2002;30(17):3894–900.

24. Wang K, Li M, Hakonarson H. ANNOVAR: functional annotation of genetic variants from high-throughput sequencing data. Nucleic Acids Res. 2010;38(16):e164.

25. Yang H, Wang K. Genomic variant annotation and prioritization with ANNOVAR and wANNOVAR. Nat Protoc. 2015;10(10):1556–66.

26. Cingolani P, Patel VM, Coon M, Nguyen T, Land SJ, Ruden DM, et al. Using Drosophila melanogaster as a Model for Genotoxic Chemical Mutational Studies with a New Program, SnpSift. Front Genet. 2012;3:35.

27. Cingolani P, Platts A, Wang le L, Coon M, Nguyen T, Wang L, et al. A program for annotating and predicting the effects of single nucleotide polymorphisms, SnpEff: SNPs in the genome of Drosophila melanogaster strain w1118; iso-2; iso-3. Fly (Austin). 2012;6(2):80-92.

28. Acharya V, Nagarajaram HA. Hansa: an automated method for discriminating disease and neutral human nsSNPs. Hum Mutat. 2012;33(2):332–7.

29. Ali H, Urolagin S, Gurarslan O, Vihinen M. Performance of protein disorder prediction programs on amino acid substitutions. Hum Mutat. 2014;35(7):794–804.

30. Alirezaie N, Kernohan KD, Hartley T, Majewski J, Hocking TD. ClinPred: Prediction Tool to Identify Disease-Relevant Nonsynonymous Single-Nucleotide Variants. Am J Hum Genet. 2018;103(4):474–83.

31. Balasubramanian S, Fu Y, Pawashe M, McGillivray P, Jin M, Liu J, et al. Using ALoFT to determine the impact of putative loss-of-function variants in protein-coding genes. Nat Commun. 2017;8(1):382.

32. Bamford S, Dawson E, Forbes S, Clements J, Pettett R, Dogan A, et al. The COSMIC (Catalogue of Somatic Mutations in Cancer) database and website. Br J Cancer. 2004;91(2):355–8.

33. Bao L, Zhou M, Cui Y. nsSNPAnalyzer: identifying disease-associated nonsynonymous single nucleotide polymorphisms. Nucleic Acids Res. 2005;33(Web Server issue):W480-2.

34. Barenboim M, Manke T. ChroMoS: an integrated web tool for SNP classification, prioritization and functional interpretation. Bioinformatics. 2013;29(17):2197–8.

35. Bendl J, Musil M, Stourac J, Zendulka J, Damborsky J, Brezovsky J. PredictSNP2: A Unified Platform for Accurately Evaluating SNP Effects by Exploiting the Different Characteristics of Variants in Distinct Genomic Regions. PLoS Comput Biol. 2016;12(5):e1004962.

36. Bendl J, Stourac J, Salanda O, Pavelka A, Wieben ED, Zendulka J, et al. PredictSNP: robust and accurate consensus classifier for prediction of disease-related mutations. PLoS Comput Biol. 2014;10(1):e1003440.

37. Bendtsen JD, Nielsen H, von Heijne G, Brunak S. Improved prediction of signal peptides: SignalP 3.0. J Mol Biol. 2004;340(4):783–95.

38. Bermejo-Das-Neves C, Nguyen HN, Poch O, Thompson JD. A comprehensive study of small non-frameshift insertions/deletions in proteins and prediction of their phenotypic effects by a machine learning method (KD4i). BMC Bioinformatics. 2014;15:111.

39. Bertoldi L, Forcato C, Vitulo N, Birolo G, De Pascale F, Feltrin E, et al. QueryOR: a comprehensive web platform for genetic variant analysis and prioritization. BMC Bioinformatics. 2017;18(1):225.

40. Bromberg Y, Rost B. SNAP: predict effect of non-synonymous polymorphisms on function. Nucleic Acids Res. 2007;35(11):3823–35.

41. Buske OJ, Manickaraj A, Mital S, Ray PN, Brudno M. Identification of deleterious synonymous variants in human genomes. Bioinformatics. 2013;29(15):1843–50.

42. Capriotti E, Altman RB. A new disease-specific machine learning approach for the prediction of cancer-causing missense variants. Genomics. 2011;98(4):310–7.

43. Capriotti E, Calabrese R, Casadio R. Predicting the insurgence of human genetic diseases associated to single point protein mutations with support vector machines and evolutionary information. Bioinformatics. 2006;22(22):2729–34.

44. Capriotti E, Calabrese R, Fariselli P, Martelli PL, Altman RB, Casadio R. WS-SNPs&GO: a web server for predicting the deleterious effect of human protein variants using functional annotation. BMC Genomics. 2013;14 Suppl 3(Suppl 3):S6.

45. Capriotti E, Casadio R. K-Fold: a tool for the prediction of the protein folding kinetic order and rate. Bioinformatics. 2007;23(3):385–6.

46. Capriotti E, Fariselli P, Calabrese R, Casadio R. Predicting protein stability changes from sequences using support vector machines. Bioinformatics. 2005;21 Suppl 2:ii54-8.

47. Cariaso M, Lennon G. SNPedia: a wiki supporting personal genome annotation, interpretation and analysis. Nucleic Acids Res. 2012;40(Database issue):D1308-12.

48. Carter H, Chen S, Isik L, Tyekucheva S, Velculescu VE, Kinzler KW, et al. Cancer-specific high-throughput annotation of somatic mutations: computational prediction of driver missense mutations. Cancer Res. 2009;69(16):6660–7.

49. Carter H, Douville C, Stenson PD, Cooper DN, Karchin R. Identifying Mendelian disease genes with the variant effect scoring tool. BMC Genomics. 2013;14 Suppl 3(Suppl 3):S3.

50. Chelala C, Khan A, Lemoine NR. SNPnexus: a web database for functional annotation of newly discovered and public domain single nucleotide polymorphisms. Bioinformatics. 2009;25(5):655–61.

51. Cheng J, Nguyen TYD, Cygan KJ, Celik MH, Fairbrother WG, Avsec Z, et al. MMSplice: modular modeling improves the predictions of genetic variant effects on splicing. Genome Biol. 2019;20(1):48.

52. Cheng J, Randall A, Baldi P. Prediction of protein stability changes for single-site mutations using support vector machines. Proteins. 2006;62(4):1125–32.

53. Choi Y, Chan AP. PROVEAN web server: a tool to predict the functional effect of amino acid substitutions and indels. Bioinformatics. 2015;31(16):2745–7.

54. Choi Y, Sims GE, Murphy S, Miller JR, Chan AP. Predicting the functional effect of amino acid substitutions and indels. PLoS One. 2012;7(10):e46688.

55. Conchillo-Sole O, de Groot NS, Aviles FX, Vendrell J, Daura X, Ventura S. AGGRESCAN: a server for the prediction and evaluation of "hot spots" of aggregation in polypeptides. BMC Bioinformatics. 2007;8:65.

56. Cuff AL, Janes RW, Martin AC. Analysing the ability to retain sidechain hydrogen-bonds in mutant proteins. Bioinformatics. 2006;22(12):1464–70.

57. Davydov EV, Goode DL, Sirota M, Cooper GM, Sidow A, Batzoglou S. Identifying a high fraction of the human genome to be under selective constraint using GERP++. PLoS Comput Biol. 2010;6(12):e1001025.

58. Dayem Ullah AZ, Lemoine NR, Chelala C. SNPnexus: a web server for functional annotation of novel and publicly known genetic variants (2012 update). Nucleic Acids Res. 2012;40(Web Server issue):W65-70.

59. Dayem Ullah AZ, Lemoine NR, Chelala C. A practical guide for the functional annotation of genetic variations using SNPnexus. Brief Bioinform. 2013;14(4):437–47.

60. Dayem Ullah AZ, Oscanoa J, Wang J, Nagano A, Lemoine NR, Chelala C. SNPnexus: assessing the functional relevance of genetic variation to facilitate the promise of precision medicine. Nucleic Acids Res. 2018;46(W1):W109–W13.

61. De Baets G, Van Durme J, Reumers J, Maurer-Stroh S, Vanhee P, Dopazo J, et al. SNPeffect 4.0: on-line prediction of molecular and structural effects of protein-coding variants. Nucleic Acids Res. 2012;40(Database issue):D935-9.

62. Dees ND, Zhang Q, Kandoth C, Wendl MC, Schierding W, Koboldt DC, et al. MuSiC: identifying mutational significance in cancer genomes. Genome Res. 2012;22(8):1589–98.

63. Dehouck Y, Kwasigroch JM, Gilis D, Rooman M. PoPMuSiC 2.1: a web server for the estimation of protein stability changes upon mutation and sequence optimality. BMC Bioinformatics. 2011;12:151.

64. Dehouck Y, Kwasigroch JM, Rooman M, Gilis D. BeAtMuSiC: Prediction of changes in protein-protein binding affinity on mutations. Nucleic Acids Res. 2013;41(Web Server issue):W333-9.

65. Desmet FO, Hamroun D, Lalande M, Collod-Beroud G, Claustres M, Beroud C. Human Splicing Finder: an online bioinformatics tool to predict splicing signals. Nucleic Acids Res. 2009;37(9):e67.

66. Deutsch C, Krishnamoorthy B. Four-body scoring function for mutagenesis. Bioinformatics. 2007;23(22):3009–15.

67. Dharanipragada P, Seelam SR, Parekh N. SeqVItA: Sequence Variant Identification and Annotation Platform for Next Generation Sequencing Data. Front Genet. 2018;9:537.

68. Dong C, Wei P, Jian X, Gibbs R, Boerwinkle E, Wang K, et al. Comparison and integration of deleteriousness prediction methods for nonsynonymous SNVs in whole exome sequencing studies. Hum Mol Genet. 2015;24(8):2125–37.

69. Dosztanyi Z, Magyar C, Tusnady G, Simon I. SCide: identification of stabilization centers in proteins. Bioinformatics. 2003;19(7):899–900.

70. Douville C, Masica DL, Stenson PD, Cooper DN, Gygax DM, Kim R, et al. Assessing the Pathogenicity of Insertion and Deletion Variants with the Variant Effect Scoring Tool (VEST-Indel). Hum Mutat. 2016;37(1):28–35.

71. Dunlavy DM, O’Leary DP, Klimov D, Thirumalai D. HOPE: a homotopy optimization method for protein structure prediction. J Comput Biol. 2005;12(10):1275-88.

72. Emanuelsson O, Brunak S, von Heijne G, Nielsen H. Locating proteins in the cell using TargetP, SignalP and related tools. Nat Protoc. 2007;2(4):953–71.

73. Fang Y, Gao S, Tai D, Middaugh CR, Fang J. Identification of properties important to protein aggregation using feature selection. BMC Bioinformatics. 2013;14:314.

74. Fariselli P, Martelli PL, Savojardo C, Casadio R. INPS: predicting the impact of non-synonymous variations on protein stability from sequence. Bioinformatics. 2015;31(17):2816–21.

75. Fernandez-Escamilla AM, Rousseau F, Schymkowitz J, Serrano L. Prediction of sequence-dependent and mutational effects on the aggregation of peptides and proteins. Nat Biotechnol. 2004;22(10):1302–6.

76. Ferrer-Costa C, Gelpi JL, Zamakola L, Parraga I, de la Cruz X, Orozco M. PMUT: a web-based tool for the annotation of pathological mutations on proteins. Bioinformatics. 2005;21(14):3176–8.

77. Fokkema IF, den Dunnen JT, Taschner PE. LOVD: easy creation of a locus-specific sequence variation database using an "LSDB-in-a-box" approach. Hum Mutat. 2005;26(2):63–8.

78. Fokkema IF, Taschner PE, Schaafsma GC, Celli J, Laros JF, den Dunnen JT. LOVD v.2.0: the next generation in gene variant databases. Hum Mutat. 2011;32(5):557–63.

79. Folkman L, Yang Y, Li Z, Stantic B, Sattar A, Mort M, et al. DDIG-in: detecting disease-causing genetic variations due to frameshifting indels and nonsense mutations employing sequence and structural properties at nucleotide and protein levels. Bioinformatics. 2015;31(10):1599–606.

80. Forbes SA, Bhamra G, Bamford S, Dawson E, Kok C, Clements J, et al. The Catalogue of Somatic Mutations in Cancer (COSMIC). Curr Protoc Hum Genet. 2008;Chapter 10:Unit 10 1.

81. Frederic MY, Lalande M, Boileau C, Hamroun D, Claustres M, Beroud C, et al. UMD-predictor, a new prediction tool for nucleotide substitution pathogenicity--application to four genes: FBN1, FBN2, TGFBR1, and TGFBR2. Hum Mutat. 2009;30(6):952-9.

82. Frousios K, Iliopoulos CS, Schlitt T, Simpson MA. Predicting the functional consequences of non-synonymous DNA sequence variants--evaluation of bioinformatics tools and development of a consensus strategy. Genomics. 2013;102(4):223–8.

83. Gao M, Skolnick J. DBD-Hunter: a knowledge-based method for the prediction of DNA-protein interactions. Nucleic Acids Res. 2008;36(12):3978–92.

84. Garber M, Guttman M, Clamp M, Zody MC, Friedman N, Xie X. Identifying novel constrained elements by exploiting biased substitution patterns. Bioinformatics. 2009;25(12):i54–62.

85. Garbuzynskiy SO, Lobanov MY, Galzitskaya OV. FoldAmyloid: a method of prediction of amyloidogenic regions from protein sequence. Bioinformatics. 2010;26(3):326–32.

86. Genomes Project C, Abecasis GR, Altshuler D, Auton A, Brooks LD, Durbin RM, et al. A map of human genome variation from population-scale sequencing. Nature. 2010;467(7319):1061-73.

87. Giollo M, Martin AJ, Walsh I, Ferrari C, Tosatto SC. NeEMO: a method using residue interaction networks to improve prediction of protein stability upon mutation. BMC Genomics. 2014;15 Suppl 4(Suppl 4):S7.

88. Goldberg T, Hamp T, Rost B. LocTree2 predicts localization for all domains of life. Bioinformatics. 2012;28(18):i458–i65.

89. Goldberg T, Hecht M, Hamp T, Karl T, Yachdav G, Ahmed N, et al. LocTree3 prediction of localization. Nucleic Acids Res. 2014;42(Web Server issue):W350-5.

90. Gonzalez-Perez A, Deu-Pons J, Lopez-Bigas N. Improving the prediction of the functional impact of cancer mutations by baseline tolerance transformation. Genome Med. 2012;4(11):89.

91. Gonzalez-Perez A, Lopez-Bigas N. Improving the assessment of the outcome of nonsynonymous SNVs with a consensus deleteriousness score, Condel. Am J Hum Genet. 2011;88(4):440–9.

92. Gosalia N, Economides AN, Dewey FE, Balasubramanian S. MAPPIN: a method for annotating, predicting pathogenicity and mode of inheritance for nonsynonymous variants. Nucleic Acids Res. 2017;45(18):10393–402.

93. Gromiha MM, Thangakani AM, Selvaraj S. FOLD-RATE: prediction of protein folding rates from amino acid sequence. Nucleic Acids Res. 2006;34(Web Server issue):W70-4.

94. Gulko B, Hubisz MJ, Gronau I, Siepel A. A method for calculating probabilities of fitness consequences for point mutations across the human genome. Nat Genet. 2015;47(3):276–83.

95. Hamosh A, Scott AF, Amberger J, Valle D, McKusick VA. Online Mendelian Inheritance in Man (OMIM). Hum Mutat. 2000;15(1):57–61.

96. Hecht M, Bromberg Y, Rost B. News from the protein mutability landscape. J Mol Biol. 2013;425(21):3937–48.

97. Hecht M, Bromberg Y, Rost B. Better prediction of functional effects for sequence variants. BMC Genomics. 2015;16 Suppl 8(Suppl 8):S1.

98. Hopf TA, Ingraham JB, Poelwijk FJ, Scharfe CP, Springer M, Sander C, et al. Mutation effects predicted from sequence co-variation. Nat Biotechnol. 2017;35(2):128–35.

99. Horton P, Park KJ, Obayashi T, Fujita N, Harada H, Adams-Collier CJ, et al. WoLF PSORT: protein localization predictor. Nucleic Acids Res. 2007;35(Web Server issue):W585-7.

100. Hu H, Huff CD, Moore B, Flygare S, Reese MG, Yandell M. VAAST 2.0: improved variant classification and disease-gene identification using a conservation-controlled amino acid substitution matrix. Genet Epidemiol. 2013;37(6):622–34.

101. Hurst JM, McMillan LE, Porter CT, Allen J, Fakorede A, Martin AC. The SAAPdb web resource: a large-scale structural analysis of mutant proteins. Hum Mutat. 2009;30(4):616–24.

102. Ioannidis NM, Rothstein JH, Pejaver V, Middha S, McDonnell SK, Baheti S, et al. REVEL: An Ensemble Method for Predicting the Pathogenicity of Rare Missense Variants. Am J Hum Genet. 2016;99(4):877–85.

103. Ionita-Laza I, McCallum K, Xu B, Buxbaum JD. A spectral approach integrating functional genomic annotations for coding and noncoding variants. Nat Genet. 2016;48(2):214–20.

104. Javed A, Agrawal S, Ng PC. Phen-Gen: combining phenotype and genotype to analyze rare disorders. Nat Methods. 2014;11(9):935–7.

105. Jia P, Zhao Z. VarWalker: personalized mutation network analysis of putative cancer genes from next-generation sequencing data. PLoS Comput Biol. 2014;10(2):e1003460.

106. Jian X, Boerwinkle E, Liu X. In silico prediction of splice-altering single nucleotide variants in the human genome. Nucleic Acids Res. 2014;42(22):13534–44.

107. Johansen MB, Izarzugaza JM, Brunak S, Petersen TN, Gupta R. Prediction of disease causing non-synonymous SNPs by the Artificial Neural Network Predictor NetDiseaseSNP. PLoS One. 2013;8(7):e68370.

108. Kaminker JS, Zhang Y, Watanabe C, Zhang Z. CanPredict: a computational tool for predicting cancer-associated missense mutations. Nucleic Acids Res. 2007;35(Web Server issue):W595-8.

109. Kang S, Chen G, Xiao G. Robust prediction of mutation-induced protein stability change by property encoding of amino acids. Protein Eng Des Sel. 2009;22(2):75–83.

110. Karczewski KJ, Francioli LC, Tiao G, Cummings BB, Alfoldi J, Wang Q, et al. The mutational constraint spectrum quantified from variation in 141,456 humans. Nature. 2020;581(7809):434-43.

111. Kircher M, Witten DM, Jain P, O’Roak BJ, Cooper GM, Shendure J. A general framework for estimating the relative pathogenicity of human genetic variants. Nat Genet. 2014;46(3):310–5.

112. Knecht C, Mort M, Junge O, Cooper DN, Krawczak M, Caliebe A. IMHOTEP-a composite score integrating popular tools for predicting the functional consequences of non-synonymous sequence variants. Nucleic Acids Res. 2017;45(3):e13.

113. Krassowski M, Paczkowska M, Cullion K, Huang T, Dzneladze I, Ouellette BFF, et al. ActiveDriverDB: human disease mutations and genome variation in post-translational modification sites of proteins. Nucleic Acids Res. 2018;46(D1):D901–D10.

114. Kulandaisamy A, Zaucha J, Sakthivel R, Frishman D, Michael Gromiha M. Pred-MutHTP: Prediction of disease-causing and neutral mutations in human transmembrane proteins. Hum Mutat. 2020;41(3):581–90.

115. Kurgan L, Cios K, Chen K. SCPRED: accurate prediction of protein structural class for sequences of twilight-zone similarity with predicting sequences. BMC Bioinformatics. 2008;9:226.

116. Laimer J, Hofer H, Fritz M, Wegenkittl S, Lackner P. MAESTRO--multi agent stability prediction upon point mutations. BMC Bioinformatics. 2015;16:116.

117. Landrum MJ, Lee JM, Benson M, Brown G, Chao C, Chitipiralla S, et al. ClinVar: public archive of interpretations of clinically relevant variants. Nucleic Acids Res. 2016;44(D1):D862–8.

118. Lappalainen I, Lopez J, Skipper L, Hefferon T, Spalding JD, Garner J, et al. DbVar and DGVa: public archives for genomic structural variation. Nucleic Acids Res. 2013;41(Database issue):D936-41.

119. Lawrence MS, Stojanov P, Polak P, Kryukov GV, Cibulskis K, Sivachenko A, et al. Mutational heterogeneity in cancer and the search for new cancer-associated genes. Nature. 2013;499(7457):214-8.

120. Lehmann KV, Chen T. Exploring functional variant discovery in non-coding regions with SInBaD. Nucleic Acids Res. 2013;41(1):e7.

121. Leiserson MD, Wu HT, Vandin F, Raphael BJ. CoMEt: a statistical approach to identify combinations of mutually exclusive alterations in cancer. Genome Biol. 2015;16(1):160.

122. Li B, Krishnan VG, Mort ME, Xin F, Kamati KK, Cooper DN, et al. Automated inference of molecular mechanisms of disease from amino acid substitutions. Bioinformatics. 2009;25(21):2744–50.

123. Li MJ, Li M, Liu Z, Yan B, Pan Z, Huang D, et al. cepip: context-dependent epigenomic weighting for prioritization of regulatory variants and disease-associated genes. Genome Biol. 2017;18(1):52.

124. Li MJ, Pan Z, Liu Z, Wu J, Wang P, Zhu Y, et al. Predicting regulatory variants with composite statistic. Bioinformatics. 2016;32(18):2729–36.

125. Li MX, Kwan JS, Bao SY, Yang W, Ho SL, Song YQ, et al. Predicting mendelian disease-causing non-synonymous single nucleotide variants in exome sequencing studies. PLoS Genet. 2013;9(1):e1003143.

126. Li Q, Wang K. InterVar: Clinical Interpretation of Genetic Variants by the 2015 ACMG-AMP Guidelines. Am J Hum Genet. 2017;100(2):267–80.

127. Linding R, Schymkowitz J, Rousseau F, Diella F, Serrano L. A comparative study of the relationship between protein structure and beta-aggregation in globular and intrinsically disordered proteins. J Mol Biol. 2004;342(1):345–53.

128. Liu M, Watson LT, Zhang L. Predicting the combined effect of multiple genetic variants. Hum Genomics. 2015;9(1):18.

129. Liu X, Jian X, Boerwinkle E. dbNSFP: a lightweight database of human nonsynonymous SNPs and their functional predictions. Hum Mutat. 2011;32(8):894–9.

130. Liu X, Jian X, Boerwinkle E. dbNSFP v2.0: a database of human non-synonymous SNVs and their functional predictions and annotations. Hum Mutat. 2013;34(9):E2393–402.

131. Liu X, White S, Peng B, Johnson AD, Brody JA, Li AH, et al. WGSA: an annotation pipeline for human genome sequencing studies. J Med Genet. 2016;53(2):111–2.

132. Liu X, Wu C, Li C, Boerwinkle E. dbNSFP v3.0: A One-Stop Database of Functional Predictions and Annotations for Human Nonsynonymous and Splice-Site SNVs. Hum Mutat. 2016;37(3):235–41.

133. Livingstone M, Folkman L, Yang Y, Zhang P, Mort M, Cooper DN, et al. Investigating DNA-, RNA-, and protein-based features as a means to discriminate pathogenic synonymous variants. Hum Mutat. 2017;38(10):1336–47.

134. Lopes MC, Joyce C, Ritchie GR, John SL, Cunningham F, Asimit J, et al. A combined functional annotation score for non-synonymous variants. Hum Hered. 2012;73(1):47–51.

135. Lopez-Ferrando V, Gazzo A, de la Cruz X, Orozco M, Gelpi JL PMut: a web-based tool for the annotation of pathological variants on proteins, 2017 update. Nucleic Acids Res. 2017;45(W1):W222-W8.

136. Lu Q, Hu Y, Sun J, Cheng Y, Cheung KH, Zhao H. A statistical framework to predict functional non-coding regions in the human genome through integrated analysis of annotation data. Sci Rep. 2015;5:10576.

137. Macintyre G, Bailey J, Haviv I, Kowalczyk A. is-rSNP: a novel technique for in silico regulatory SNP detection. Bioinformatics. 2010;26(18):i524–30.

138. Mao Y, Chen H, Liang H, Meric-Bernstam F, Mills GB, Chen K. CanDrA: cancer-specific driver missense mutation annotation with optimized features. PLoS One. 2013;8(10):e77945.

139. Marini NJ, Thomas PD, Rine J. The use of orthologous sequences to predict the impact of amino acid substitutions on protein function. PLoS Genet. 2010;6(5):e1000968.

140. Masso M, Vaisman, II. AUTO-MUTE: web-based tools for predicting stability changes in proteins due to single amino acid replacements. Protein Eng Des Sel. 2010;23(8):683–7.

141. Mathe E, Olivier M, Kato S, Ishioka C, Hainaut P, Tavtigian SV. Computational approaches for predicting the biological effect of p53 missense mutations: a comparison of three sequence analysis based methods. Nucleic Acids Res. 2006;34(5):1317–25.

142. Maurer-Stroh S, Debulpaep M, Kuemmerer N, Lopez de la Paz M, Martins IC, Reumers J, et al. Exploring the sequence determinants of amyloid structure using position-specific scoring matrices. Nat Methods. 2010;7(3):237–42.

143. McLaren W, Gil L, Hunt SE, Riat HS, Ritchie GR, Thormann A, et al. The Ensembl Variant Effect Predictor. Genome Biol. 2016;17(1):122.

144. McLaren W, Pritchard B, Rios D, Chen Y, Flicek P, Cunningham F. Deriving the consequences of genomic variants with the Ensembl API and SNP Effect Predictor. Bioinformatics. 2010;26(16):2069–70.

145. Mi H, Huang X, Muruganujan A, Tang H, Mills C, Kang D, et al. PANTHER version 11: expanded annotation data from Gene Ontology and Reactome pathways, and data analysis tool enhancements. Nucleic Acids Res. 2017;45(D1):D183–D9.

146. Moretti R, Fleishman SJ, Agius R, Torchala M, Bates PA, Kastritis PL, et al. Community-wide evaluation of methods for predicting the effect of mutations on protein-protein interactions. Proteins. 2013;81(11):1980–7.

147. Mort M, Sterne-Weiler T, Li B, Ball EV, Cooper DN, Radivojac P, et al. MutPred Splice: machine learning-based prediction of exonic variants that disrupt splicing. Genome Biol. 2014;15(1):R19.

148. Nalla VK, Rogan PK. Automated splicing mutation analysis by information theory. Hum Mutat. 2005;25(4):334–42.

149. Nielsen H, Krogh A. Prediction of signal peptides and signal anchors by a hidden Markov model. Proc Int Conf Intell Syst Mol Biol. 1998;6:122–30.

150. Niroula A, Urolagin S, Vihinen M. PON-P2: prediction method for fast and reliable identification of harmful variants. PLoS One. 2015;10(2):e0117380.

151. Niroula A, Vihinen M. PON-mt-tRNA: a multifactorial probability-based method for classification of mitochondrial tRNA variations. Nucleic Acids Res. 2016;44(5):2020–7.

152. Olatubosun A, Valiaho J, Harkonen J, Thusberg J, Vihinen M. PON-P: integrated predictor for pathogenicity of missense variants. Hum Mutat. 2012;33(8):1166–74.

153. Pagel KA, Pejaver V, Lin GN, Nam HJ, Mort M, Cooper DN, et al. When loss-of-function is loss of function: assessing mutational signatures and impact of loss-of-function genetic variants. Bioinformatics. 2017;33(14):i389–i98.

154. Pagon RA, Tarczy-Hornoch P, Baskin PK, Edwards JE, Covington ML, Espeseth M, et al. GeneTests-GeneClinics: genetic testing information for a growing audience. Hum Mutat. 2002;19(5):501–9.

155. Pandurangan AP, Ochoa-Montano B, Ascher DB, Blundell TL. SDM: a server for predicting effects of mutations on protein stability. Nucleic Acids Res. 2017;45(W1):W229–W35.

156. Pappalardo M, Wass MN. VarMod: modelling the functional effects of non-synonymous variants. Nucleic Acids Res. 2014;42(Web Server issue):W331-6.

157. Parthiban V, Gromiha MM, Abhinandan M, Schomburg D. Computational modeling of protein mutant stability: analysis and optimization of statistical potentials and structural features reveal insights into prediction model development. BMC Struct Biol. 2007;7:54.

158. Parthiban V, Gromiha MM, Hoppe C, Schomburg D. Structural analysis and prediction of protein mutant stability using distance and torsion potentials: role of secondary structure and solvent accessibility. Proteins. 2007;66(1):41–52.

159. Parthiban V, Gromiha MM, Schomburg D. CUPSAT: prediction of protein stability upon point mutations. Nucleic Acids Res. 2006;34(Web Server issue):W239-42.

160. Pejaver V, Urresti J, Lugo-Martinez J, Pagel KA, Lin GN, Nam HJ, et al. Inferring the molecular and phenotypic impact of amino acid variants with MutPred2. Nat Commun. 2020;11(1):5918.

161. Peng B. Reproducible simulations of realistic samples for next-generation sequencing studies using Variant Simulation Tools. Genet Epidemiol. 2015;39(1):45–52.

162. Petersen TN, Brunak S, von Heijne G, Nielsen H. SignalP 4.0: discriminating signal peptides from transmembrane regions. Nat Methods. 2011;8(10):785–6.

163. Pires DE, Ascher DB, Blundell TL. DUET: a server for predicting effects of mutations on protein stability using an integrated computational approach. Nucleic Acids Res. 2014;42(Web Server issue):W314-9.

164. Pokala N, Handel TM. Energy functions for protein design: adjustment with protein-protein complex affinities, models for the unfolded state, and negative design of solubility and specificity. J Mol Biol. 2005;347(1):203–27.

165. Pollard KS, Hubisz MJ, Rosenbloom KR, Siepel A. Detection of nonneutral substitution rates on mammalian phylogenies. Genome Res. 2010;20(1):110–21.

166. Preeprem T, Gibson G. SDS, a structural disruption score for assessment of missense variant deleteriousness. Front Genet. 2014;5:82.

167. Punta M, Rost B. PROFcon: novel prediction of long-range contacts. Bioinformatics. 2005;21(13):2960–8.

168. Qin S, Pang X, Zhou HX. Automated prediction of protein association rate constants. Structure. 2011;19(12):1744–51.

169. Quang D, Chen Y, Xie X. DANN: a deep learning approach for annotating the pathogenicity of genetic variants. Bioinformatics. 2015;31(5):761–3.

170. Reumers J, Conde L, Medina I, Maurer-Stroh S, Van Durme J, Dopazo J, et al. Joint annotation of coding and non-coding single nucleotide polymorphisms and mutations in the SNPeffect and PupaSuite databases. Nucleic Acids Res. 2008;36(Database issue):D825-9.

171. Reumers J, Maurer-Stroh S, Schymkowitz J, Rousseau F. SNPeffect v2.0: a new step in investigating the molecular phenotypic effects of human non-synonymous SNPs. Bioinformatics. 2006;22(17):2183–5.

172. Reumers J, Schymkowitz J, Ferkinghoff-Borg J, Stricher F, Serrano L, Rousseau F. SNPeffect: a database mapping molecular phenotypic effects of human non-synonymous coding SNPs. Nucleic Acids Res. 2005;33(Database issue):D527-32.

173. Reva B, Antipin Y, Sander C. Determinants of protein function revealed by combinatorial entropy optimization. Genome Biol. 2007;8(11):R232.

174. Reva B, Antipin Y, Sander C. Predicting the functional impact of protein mutations: application to cancer genomics. Nucleic Acids Res. 2011;39(17):e118.

175. Ritchie GR, Dunham I, Zeggini E, Flicek P. Functional annotation of noncoding sequence variants. Nat Methods. 2014;11(3):294–6.

176. Rousseau F, Schymkowitz J, Serrano L. Protein aggregation and amyloidosis: confusion of the kinds? Curr Opin Struct Biol. 2006;16(1):118–26.

177. Ryan M, Diekhans M, Lien S, Liu Y, Karchin R. LS-SNP/PDB: annotated non-synonymous SNPs mapped to Protein Data Bank structures. Bioinformatics. 2009;25(11):1431–2.

178. Ryan NM, Morris SW, Porteous DJ, Taylor MS, Evans KL. SuRFing the genomics wave: an R package for prioritising SNPs by functionality. Genome Med. 2014;6(10):79.

179. San Lucas FA, Wang G, Scheet P, Peng B. Integrated annotation and analysis of genetic variants from next-generation sequencing studies with variant tools. Bioinformatics. 2012;28(3):421–2.

180. Sasidharan Nair P, Vihinen M. VariBench: a benchmark database for variations. Hum Mutat. 2013;34(1):42–9.

181. Savojardo C, Fariselli P, Martelli PL, Casadio R. INPS-MD: a web server to predict stability of protein variants from sequence and structure. Bioinformatics. 2016;32(16):2542–4.

182. Schaafsma GC, Vihinen M. VariSNP, a benchmark database for variations from dbSNP. Hum Mutat. 2015;36(2):161–6.

183. Schaefer C, Meier A, Rost B, Bromberg Y. SNPdbe: constructing an nsSNP functional impacts database. Bioinformatics. 2012;28(4):601–2.

184. Schwarz JM, Cooper DN, Schuelke M, Seelow D. MutationTaster2: mutation prediction for the deep-sequencing age. Nat Methods. 2014;11(4):361–2.

185. Schwarz JM, Rodelsperger C, Schuelke M, Seelow D. MutationTaster evaluates disease-causing potential of sequence alterations. Nat Methods. 2010;7(8):575–6.

186. Schymkowitz J, Borg J, Stricher F, Nys R, Rousseau F, Serrano L. The FoldX web server: an online force field. Nucleic Acids Res. 2005;33(Web Server issue):W382-8.

187. Sherry ST, Ward MH, Kholodov M, Baker J, Phan L, Smigielski EM, et al. dbSNP: the NCBI database of genetic variation. Nucleic Acids Res. 2001;29(1):308–11.

188. Shihab HA, Gough J, Cooper DN, Day IN, Gaunt TR. Predicting the functional consequences of cancer-associated amino acid substitutions. Bioinformatics. 2013;29(12):1504–10.

189. Shihab HA, Gough J, Cooper DN, Stenson PD, Barker GL, Edwards KJ, et al. Predicting the functional, molecular, and phenotypic consequences of amino acid substitutions using hidden Markov models. Hum Mutat. 2013;34(1):57–65.

190. Shihab HA, Gough J, Mort M, Cooper DN, Day IN, Gaunt TR. Ranking non-synonymous single nucleotide polymorphisms based on disease concepts. Hum Genomics. 2014;8(1):11.

191. Shihab HA, Rogers MF, Gough J, Mort M, Cooper DN, Day IN, et al. An integrative approach to predicting the functional effects of non-coding and coding sequence variation. Bioinformatics. 2015;31(10):1536–43.

192. Shringarpure SS, Bustamante CD. Privacy Risks from Genomic Data-Sharing Beacons. Am J Hum Genet. 2015;97(5):631–46.

193. Siepel A, Bejerano G, Pedersen JS, Hinrichs AS, Hou M, Rosenbloom K, et al. Evolutionarily conserved elements in vertebrate, insect, worm, and yeast genomes. Genome Res. 2005;15(8):1034–50.

194. Smedley D, Jacobsen JO, Jager M, Kohler S, Holtgrewe M, Schubach M, et al. Next-generation diagnostics and disease-gene discovery with the Exomiser. Nat Protoc. 2015;10(12):2004–15.

195. Smedley D, Schubach M, Jacobsen JOB, Kohler S, Zemojtel T, Spielmann M, et al. A Whole-Genome Analysis Framework for Effective Identification of Pathogenic Regulatory Variants in Mendelian Disease. Am J Hum Genet. 2016;99(3):595–606.

196. Stenson PD, Mort M, Ball EV, Evans K, Hayden M, Heywood S, et al. The Human Gene Mutation Database: towards a comprehensive repository of inherited mutation data for medical research, genetic diagnosis and next-generation sequencing studies. Hum Genet. 2017;136(6):665–77.

197. Stone EA, Sidow A. Physicochemical constraint violation by missense substitutions mediates impairment of protein function and disease severity. Genome Res. 2005;15(7):978–86.

198. Tamborero D, Gonzalez-Perez A, Lopez-Bigas N. OncodriveCLUST: exploiting the positional clustering of somatic mutations to identify cancer genes. Bioinformatics. 2013;29(18):2238–44.

199. Tang H, Thomas PD. PANTHER-PSEP: predicting disease-causing genetic variants using position-specific evolutionary preservation. Bioinformatics. 2016;32(14):2230–2.

200. Tartaglia GG, Cavalli A, Pellarin R, Caflisch A. Prediction of aggregation rate and aggregation-prone segments in polypeptide sequences. Protein Sci. 2005;14(10):2723–34.

201. Tartaglia GG, Vendruscolo M. The Zyggregator method for predicting protein aggregation propensities. Chem Soc Rev. 2008;37(7):1395–401.

202. Tate JG, Bamford S, Jubb HC, Sondka Z, Beare DM, Bindal N, et al. COSMIC: the Catalogue Of Somatic Mutations In Cancer. Nucleic Acids Res. 2019;47(D1):D941–D7.

203. Tavtigian SV, Deffenbaugh AM, Yin L, Judkins T, Scholl T, Samollow PB, et al. Comprehensive statistical study of 452 BRCA1 missense substitutions with classification of eight recurrent substitutions as neutral. J Med Genet. 2006;43(4):295–305.

204. Teng S, Srivastava AK, Wang L. Sequence feature-based prediction of protein stability changes upon amino acid substitutions. BMC Genomics. 2010;11 Suppl 2(Suppl 2):S5.

205. Terui H, Akagi K, Kawame H, Yura K. CoDP: predicting the impact of unclassified genetic variants in MSH6 by the combination of different properties of the protein. J Biomed Sci. 2013;20(1):25.

206. Thompson BA, Spurdle AB, Plazzer JP, Greenblatt MS, Akagi K, Al-Mulla F, et al. Application of a 5-tiered scheme for standardized classification of 2,360 unique mismatch repair gene variants in the InSiGHT locus-specific database. Nat Genet. 2014;46(2):107–15.

207. Thorn CF, Klein TE, Altman RB. PharmGKB: the Pharmacogenomics Knowledge Base. Methods Mol Biol. 2013;1015:311–20.

208. Tian J, Wu N, Guo X, Guo J, Zhang J, Fan Y. Predicting the phenotypic effects of non-synonymous single nucleotide polymorphisms based on support vector machines. BMC Bioinformatics. 2007;8:450.

209. Vuong H, Che A, Ravichandran S, Luke BT, Collins JR, Mudunuri US. AVIA v2.0: annotation, visualization and impact analysis of genomic variants and genes. Bioinformatics. 2015;31(16):2748–50.

210. Walsh I, Seno F, Tosatto SC, Trovato A. PASTA 2.0: an improved server for protein aggregation prediction. Nucleic Acids Res. 2014;42(Web Server issue):W301-7.

211. Wang GT, Peng B, Leal SM. Variant association tools for quality control and analysis of large-scale sequence and genotyping array data. Am J Hum Genet. 2014;94(5):770–83.

212. Wang M, Zhao XM, Takemoto K, Xu H, Li Y, Akutsu T, et al. FunSAV: predicting the functional effect of single amino acid variants using a two-stage random forest model. PLoS One. 2012;7(8):e43847.

213. Wishart DS, Arndt D, Berjanskii M, Guo AC, Shi Y, Shrivastava S, et al. PPT-DB: the protein property prediction and testing database. Nucleic Acids Res. 2008;36(Database issue):D222-9.

214. Wong WC, Kim D, Carter H, Diekhans M, Ryan MC, Karchin R. CHASM and SNVBox: toolkit for detecting biologically important single nucleotide mutations in cancer. Bioinformatics. 2011;27(15):2147–8.

215. Woolfe A, Mullikin JC, Elnitski L. Genomic features defining exonic variants that modulate splicing. Genome Biol. 2010;11(2):R20.

216. Xiong HY, Alipanahi B, Lee LJ, Bretschneider H, Merico D, Yuen RK, et al. RNA splicing. The human splicing code reveals new insights into the genetic determinants of disease. Science. 2015;347(6218):1254806.

217. Xu B, Yang Y, Liang H, Zhou Y. An all-atom knowledge-based energy function for protein-DNA threading, docking decoy discrimination, and prediction of transcription-factor binding profiles. Proteins. 2009;76(3):718–30.

218. Xu Z, Taylor JA. SNPinfo: integrating GWAS and candidate gene information into functional SNP selection for genetic association studies. Nucleic Acids Res. 2009;37(Web Server issue):W600-5.

219. Yandell M, Huff C, Hu H, Singleton M, Moore B, Xing J, et al. A probabilistic disease-gene finder for personal genomes. Genome Res. 2011;21(9):1529–42.

220. Ye ZQ, Zhao SQ, Gao G, Liu XQ, Langlois RE, Lu H, et al. Finding new structural and sequence attributes to predict possible disease association of single amino acid polymorphism (SAP). Bioinformatics. 2007;23(12):1444–50.

221. Yeo G, Burge CB. Maximum entropy modeling of short sequence motifs with applications to RNA splicing signals. J Comput Biol. 2004;11(2-3):377–94.

222. Yin S, Ding F, Dokholyan NV. Modeling backbone flexibility improves protein stability estimation. Structure. 2007;15(12):1567–76.

223. Yin S, Ding F, Dokholyan NV. Eris: an automated estimator of protein stability. Nat Methods. 2007;4(6):466–7.

224. Yue P, Melamud E, Moult J. SNPs3D: candidate gene and SNP selection for association studies. BMC Bioinformatics. 2006;7:166.

225. Yue P, Moult J. Identification and analysis of deleterious human SNPs. J Mol Biol. 2006;356(5):1263–74.

226. Zambrano R, Jamroz M, Szczasiuk A, Pujols J, Kmiecik S, Ventura S. AGGRESCAN3D (A3D): server for prediction of aggregation properties of protein structures. Nucleic Acids Res. 2015;43(W1):W306–13.

227. Zeng S, Yang J, Chung BH, Lau YL, Yang W. EFIN: predicting the functional impact of nonsynonymous single nucleotide polymorphisms in human genome. BMC Genomics. 2014;15(1):455.

228. Zhang C, Liu S, Zhu Q, Zhou Y. A knowledge-based energy function for protein-ligand, protein-protein, and protein-DNA complexes. J Med Chem. 2005;48(7):2325–35.

229. Zhang T, Wu Y, Lan Z, Shi Q, Yang Y, Guo J. Syntool: A Novel Region-Based Intolerance Score to Single Nucleotide Substitution for Synonymous Mutations Predictions Based on 123,136 Individuals. Biomed Res Int. 2017;2017:5096208.

230. Zhao H, Yang Y, Lin H, Zhang X, Mort M, Cooper DN, et al. DDIG-in: discriminating between disease-associated and neutral non-frameshifting micro-indels. Genome Biol. 2013;14(3):R23.

231. Zhou H, Zhou Y. Distance-scaled, finite ideal-gas reference state improves structure-derived potentials of mean force for structure selection and stability prediction. Protein Sci. 2002;11(11):2714–26.

232. Zhou J, Theesfeld CL, Yao K, Chen KM, Wong AK, Troyanskaya OG. Deep learning sequence-based ab initio prediction of variant effects on expression and disease risk. Nat Genet. 2018;50(8):1171–9.

233. Zhou J, Troyanskaya OG. Predicting effects of noncoding variants with deep learning-based sequence model. Nat Methods. 2015;12(10):931–4.

234. Addepalli A, Kalyani S, Singh M, Bandyopadhyay D, Mohan KN. CalPen (Calculator of Penetrance), a web-based tool to estimate penetrance in complex genetic disorders. PLoS One. 2020;15(1):e0228156.

235. Alexander J, Mantzaris D, Georgitsi M, Drineas P, Paschou P. Variant Ranker: a web-tool to rank genomic data according to functional significance. BMC Bioinformatics. 2017;18(1):341.

236. Allot A, Peng Y, Wei CH, Lee K, Phan L, Lu Z. LitVar: a semantic search engine for linking genomic variant data in PubMed and PMC. Nucleic Acids Res. 2018;46(W1):W530–W6.

237. Ancien F, Pucci F, Godfroid M, Rooman M. Prediction and interpretation of deleterious coding variants in terms of protein structural stability. Sci Rep. 2018;8(1):4480.

238. Arani AA, Sehhati M, Tabatabaiefar MA. Genetic variant effect prediction by supervised nonnegative matrix tri-factorization. Mol Omics. 2021;17(5):740–51.

239. Avsec Z, Agarwal V, Visentin D, Ledsam JR, Grabska-Barwinska A, Taylor KR, et al. Effective gene expression prediction from sequence by integrating long-range interactions. Nat Methods. 2021;18(10):1196–203.

240. Bailey M, Miller N. DMD Open-access Variant Explorer (DOVE): A scalable, open-access, web-based tool to aid in clinical interpretation of genetic variants in the DMD gene. Mol Genet Genomic Med. 2019;7(1):e00510.

241. Barbon L, Offord V, Radford EJ, Butler AP, Gerety SS, Adams DJ, et al. Variant Library Annotation Tool (VaLiAnT): an oligonucleotide library design and annotation tool for saturation genome editing and other deep mutational scanning experiments. Bioinformatics. 2022;38(4):892–9.

242. Basile AO, Byrska-Bishop M, Wallace J, Frase AT, Ritchie MD. Novel features and enhancements in BioBin, a tool for the biologically inspired binning and association analysis of rare variants. Bioinformatics. 2018;34(3):527–9.

243. Benegas G, Batra SS, Song YS. DNA language models are powerful predictors of genome-wide variant effects. Proc Natl Acad Sci U S A. 2023;120(44):e2311219120.

244. Benton MC, Smith RA, Haupt LM, Sutherland HG, Dunn PJ, Albury CL, et al. Variant Call Format-Diagnostic Annotation and Reporting Tool: A Customizable Analysis Pipeline for Identification of Clinically Relevant Genetic Variants in Next-Generation Sequencing Data. J Mol Diagn. 2019;21(6):951–60.

245. Bhattacharya S, Barseghyan H, Delot EC, Vilain E. nanotatoR: a tool for enhanced annotation of genomic structural variants. BMC Genomics. 2021;22(1):10.

246. Binatti A, Bresolin S, Bortoluzzi S, Coppe A. iWhale: a computational pipeline based on Docker and SCons for detection and annotation of somatic variants in cancer WES data. Brief Bioinform. 2021;22(3).

247. Buniello A, MacArthur JAL, Cerezo M, Harris LW, Hayhurst J, Malangone C, et al. The NHGRI-EBI GWAS Catalog of published genome-wide association studies, targeted arrays and summary statistics 2019. Nucleic Acids Res. 2019;47(D1):D1005–D12.

248. Calabrese R, Capriotti E, Fariselli P, Martelli PL, Casadio R. Functional annotations improve the predictive score of human disease-related mutations in proteins. Hum Mutat. 2009;30(8):1237–44.

249. Cao H, Wang J, He L, Qi Y, Zhang JZ. DeepDDG: Predicting the Stability Change of Protein Point Mutations Using Neural Networks. J Chem Inf Model. 2019;59(4):1508–14.

250. Cao Y, Ha SY, So CC, Tong MT, Tang CS, Zhang H, et al. NGS4THAL, a One-Stop Molecular Diagnosis and Carrier Screening Tool for Thalassemia and Other Hemoglobinopathies by Next-Generation Sequencing. J Mol Diagn. 2022;24(10):1089–99.

251. Capriotti E, Fariselli P. PhD-SNPg: updating a webserver and lightweight tool for scoring nucleotide variants. Nucleic Acids Res. 2023;51(W1):W451–W8.

252. Chakravarty D, Gao J, Phillips SM, Kundra R, Zhang H, Wang J, et al. OncoKB: A Precision Oncology Knowledge Base. JCO Precis Oncol. 2017;2017.

253. Chang MT, Bhattarai TS, Schram AM, Bielski CM, Donoghue MTA, Jonsson P, et al. Accelerating Discovery of Functional Mutant Alleles in Cancer. Cancer Discov. 2018;8(2):174–83.

254. Chen CW, Lin J, Chu YW. iStable: off-the-shelf predictor integration for predicting protein stability changes. BMC Bioinformatics. 2013;14 Suppl 2(Suppl 2):S5.

255. Chen CW, Lin MH, Liao CC, Chang HP, Chu YW. iStable 2.0: Predicting protein thermal stability changes by integrating various characteristic modules. Comput Struct Biotechnol J. 2020;18:622–30.

256. Chen Y, Lu H, Zhang N, Zhu Z, Wang S, Li M. PremPS: Predicting the impact of missense mutations on protein stability. PLoS Comput Biol. 2020;16(12):e1008543.

257. Cheng J, Novati G, Pan J, Bycroft C, Zemgulyte A, Applebaum T, et al. Accurate proteome-wide missense variant effect prediction with AlphaMissense. Science. 2023;381(6664):eadg7492.

258. Chennen K, Weber T, Lornage X, Kress A, Bohm J, Thompson J, et al. MISTIC: A prediction tool to reveal disease-relevant deleterious missense variants. PLoS One. 2020;15(7):e0236962.

259. Chun S, Fay JC. Identification of deleterious mutations within three human genomes. Genome Res. 2009;19(9):1553–61.

260. Cipriani V, Pontikos N, Arno G, Sergouniotis PI, Lenassi E, Thawong P, et al. An Improved Phenotype-Driven Tool for Rare Mendelian Variant Prioritization: Benchmarking Exomiser on Real Patient Whole-Exome Data. Genes (Basel). 2020;11(4).

261. Clausen R, Ma B, Nussinov R, Shehu A. Mapping the Conformation Space of Wildtype and Mutant H-Ras with a Memetic, Cellular, and Multiscale Evolutionary Algorithm. PLoS Comput Biol. 2015;11(9):e1004470.

262. Cooper DN, Ball EV, Krawczak M. The human gene mutation database. Nucleic Acids Res. 1998;26(1):285–7.

263. Cooper DN, Stenson PD, Chuzhanova NA. The Human Gene Mutation Database (HGMD) and its exploitation in the study of mutational mechanisms. Curr Protoc Bioinformatics. 2006;Chapter 1:Unit 1 13.

264. Costanzo MC, Roselli C, Brandes M, Duby M, Hoang Q, Jang D, et al. Cardiovascular Disease Knowledge Portal: A Community Resource for Cardiovascular Disease Research. Circ Genom Precis Med. 2023;16(6):e004181.

265. Danis D, Jacobsen JOB, Balachandran P, Zhu Q, Yilmaz F, Reese J, et al. SvAnna: efficient and accurate pathogenicity prediction of coding and regulatory structural variants in long-read genome sequencing. Genome Med. 2022;14(1):44.

266. Danis D, Jacobsen JOB, Carmody LC, Gargano MA, McMurry JA, Hegde A, et al. Interpretable prioritization of splice variants in diagnostic next-generation sequencing. Am J Hum Genet. 2021;108(9):1564–77.

267. Danzi MC, Dohrn MF, Fazal S, Beijer D, Rebelo AP, Cintra V, et al. Deep structured learning for variant prioritization in Mendelian diseases. Nat Commun. 2023;14(1):4167.

268. Derbel H, Zhao Z, Liu Q. Accurate prediction of functional effect of single amino acid variants with deep learning. Comput Struct Biotechnol J. 2023;21:5776–84.

269. Di Sera T, Velinder M, Ward A, Qiao Y, Georges S, Miller C, et al. Gene.iobio: an interactive web tool for versatile, clinically-driven variant interrogation and prioritization. Sci Rep. 2021;11(1):20307.

270. Dunham AS, Beltrao P, AlQuraishi M. High-throughput deep learning variant effect prediction with Sequence UNET. Genome Biol. 2023;24(1):110.

271. Ekawade A, Velinder M, Ward A, DiSera T, Miller C, Qiao Y, et al. Genepanel.iobio-an easy to use web tool for generating disease-and phenotype-associated gene lists. BMC Med Genomics. 2019;12(1):190.

272. Esposito D, Weile J, Shendure J, Starita LM, Papenfuss AT, Roth FP, et al. MaveDB: an open-source platform to distribute and interpret data from multiplexed assays of variant effect. Genome Biol. 2019;20(1):223.

273. Fang M, Su Z, Abolhassani H, Itan Y, Jin X, Hammarstrom L. VIPPID: a gene-specific single nucleotide variant pathogenicity prediction tool for primary immunodeficiency diseases. Brief Bioinform. 2022;23(5).

274. Feng BJ. PERCH: A Unified Framework for Disease Gene Prioritization. Hum Mutat. 2017;38(3):243–51.

275. Frazer J, Notin P, Dias M, Gomez A, Min JK, Brock K, et al. Disease variant prediction with deep generative models of evolutionary data. Nature. 2021;599(7883):91-5.

276. Fredrich B, Schmohl M, Junge O, Gundlach S, Ellinghaus D, Pfeufer A, et al. VarWatch-A stand-alone software tool for variant matching. PLoS One. 2019;14(4):e0215618.

277. Fu Y, Liu Z, Lou S, Bedford J, Mu XJ, Yip KY, et al. FunSeq2: a framework for prioritizing noncoding regulatory variants in cancer. Genome Biol. 2014;15(10):480.

278. Galano-Frutos JJ, Garcia-Cebollada H, Lopez A, Rosell M, de la Cruz X, Fernandez-Recio J, et al. PirePred: An Accurate Online Consensus Tool to Interpret Newborn Screening-Related Genetic Variants in Structural Context. J Mol Diagn. 2022;24(4):406–25.

279. Ganel L, Abel HJ, FinMetSeq C, Hall IM. SVScore: an impact prediction tool for structural variation. Bioinformatics. 2017;33(7):1083–5.

280. Ganesan K, Kulandaisamy A, Binny Priya S, Gromiha MM. HuVarBase: A human variant database with comprehensive information at gene and protein levels. PLoS One. 2019;14(1):e0210475.

281. Gao H, Hamp T, Ede J, Schraiber JG, McRae J, Singer-Berk M, et al. The landscape of tolerated genetic variation in humans and primates. Science. 2023;380(6648):eabn8153.

282. Gazzo AM, Daneels D, Cilia E, Bonduelle M, Abramowicz M, Van Dooren S, et al. DIDA: A curated and annotated digenic diseases database. Nucleic Acids Res. 2016;44(D1):D900–7.

283. Geoffroy V, Pizot C, Redin C, Piton A, Vasli N, Stoetzel C, et al. VaRank: a simple and powerful tool for ranking genetic variants. PeerJ. 2015;3:e796.

284. Glanzmann B, Herbst H, Kinnear CJ, Moller M, Gamieldien J, Bardien S. A new tool for prioritization of sequence variants from whole exome sequencing data. Source Code Biol Med. 2016;11:10.

285. Gong J, Wang J, Zong X, Ma Z, Xu D. Prediction of protein stability changes upon single-point variant using 3D structure profile. Comput Struct Biotechnol J. 2023;21:354–64.

286. Granata I, Sangiovanni M, Maiorano F, Miele M, Guarracino MR. Var2GO: a web-based tool for gene variants selection. BMC Bioinformatics. 2016;17(Suppl 12):376.

287. Griffith M, Spies NC, Krysiak K, McMichael JF, Coffman AC, Danos AM, et al. CIViC is a community knowledgebase for expert crowdsourcing the clinical interpretation of variants in cancer. Nat Genet. 2017;49(2):170–4.

288. Guo Y, Tian K, Zeng H, Guo X, Gifford DK. A novel k-mer set memory (KSM) motif representation improves regulatory variant prediction. Genome Res. 2018;28(6):891–900.

289. Gurbich TA, Ilinsky VV. ClassifyCNV: a tool for clinical annotation of copy-number variants. Sci Rep. 2020;10(1):20375.

290. Han Q, Yang Y, Wu S, Liao Y, Zhang S, Liang H, et al. Cruxome: a powerful tool for annotating, interpreting and reporting genetic variants. BMC Genomics. 2021;22(1):407.

291. Hart SN, Polley EC, Shimelis H, Yadav S, Couch FJ. Prediction of the functional impact of missense variants in BRCA1 and BRCA2 with BRCA-ML. NPJ Breast Cancer. 2020;6:13.

292. He MM, Li Q, Yan M, Cao H, Hu Y, He KY, et al. Variant Interpretation for Cancer (VIC): a computational tool for assessing clinical impacts of somatic variants. Genome Med. 2019;11(1):53.

293. Howard M, Kane B, Lepry M, Stey P, Ragavendran A, Gamsiz Uzun ED. VarStack: a web tool for data retrieval to interpret somatic variants in cancer. Database (Oxford). 2020;2020.

294. Huang YF, Gulko B, Siepel A. Fast, scalable prediction of deleterious noncoding variants from functional and population genomic data. Nat Genet. 2017;49(4):618–24.

295. Ip E, Chapman G, Winlaw D, Dunwoodie SL, Giannoulatou E. VPOT: A Customizable Variant Prioritization Ordering Tool for Annotated Variants. Genomics Proteomics Bioinformatics. 2019;17(5):540–5.

296. Iqbal S, Hoksza D, Perez-Palma E, May P, Jespersen JB, Ahmed SS, et al. MISCAST: MIssense variant to protein StruCture Analysis web SuiTe. Nucleic Acids Res. 2020;48(W1):W132–W9.

297. Jagadeesh KA, Wenger AM, Berger MJ, Guturu H, Stenson PD, Cooper DN, et al. M-CAP eliminates a majority of variants of uncertain significance in clinical exomes at high sensitivity. Nat Genet. 2016;48(12):1581–6.

298. Jaganathan K, Kyriazopoulou Panagiotopoulou S, McRae JF, Darbandi SF, Knowles D, Li YI, et al. Predicting Splicing from Primary Sequence with Deep Learning. Cell. 2019;176(3):535–48 e24.

299. Jagota M, Ye C, Albors C, Rastogi R, Koehl A, Ioannidis N, et al. Cross-protein transfer learning substantially improves disease variant prediction. Genome Biol. 2023;24(1):182.

300. Jiang S, Xie Y, He Z, Zhang Y, Zhao Y, Chen L, et al. m6ASNP: a tool for annotating genetic variants by m6A function. Gigascience. 2018;7(5).

301. Kaakinen M, Magi R, Fischer K, Heikkinen J, Jarvelin MR, Morris AP, et al. MARV: a tool for genome-wide multi-phenotype analysis of rare variants. BMC Bioinformatics. 2017;18(1):110.

302. Kalayci S, Selvan ME, Ramos I, Cotsapas C, Harris E, Kim EY, et al. ImmuneRegulation: a web-based tool for identifying human immune regulatory elements. Nucleic Acids Res. 2019;47(W1):W142–W50.

303. Kamat MA, Blackshaw JA, Young R, Surendran P, Burgess S, Danesh J, et al. PhenoScanner V2: an expanded tool for searching human genotype-phenotype associations. Bioinformatics. 2019;35(22):4851–3.

304. Karmakar M, Cicaloni V, Rodrigues CHM, Spiga O, Santucci A, Ascher DB. HGDiscovery: An online tool providing functional and phenotypic information on novel variants of homogentisate 1,2-dioxigenase. Curr Res Struct Biol. 2022;4:271–7.

305. Kasaragod S, Mohanty V, Tyagi A, Behera SK, Patil AH, Pinto SM, et al. CusVarDB: A tool for building customized sample-specific variant protein database from next-generation sequencing datasets. F1000Res. 2020;9:344.

306. Katsonis P, Lichtarge O. A formal perturbation equation between genotype and phenotype determines the Evolutionary Action of protein-coding variations on fitness. Genome Res. 2014;24(12):2050–8.

307. Krawczak M, Ball EV, Fenton I, Stenson PD, Abeysinghe S, Thomas N, et al. Human gene mutation database-a biomedical information and research resource. Hum Mutat. 2000;15(1):45–51.

308. Kulandaisamy A, Binny Priya S, Sakthivel R, Tarnovskaya S, Bizin I, Honigschmid P, et al. MutHTP: mutations in human transmembrane proteins. Bioinformatics. 2018;34(13):2325–6.

309. Kulandaisamy A, Zaucha J, Frishman D, Gromiha MM. MPTherm-pred: Analysis and Prediction of Thermal Stability Changes upon Mutations in Transmembrane Proteins. J Mol Biol. 2021;433(11):166646.

310. Laddach A, Gautel M, Fraternali F. TITINdb-a computational tool to assess titin’s role as a disease gene. Bioinformatics. 2017;33(21):3482–5.

311. Lai C, Zimmer AD, O’Connor R, Kim S, Chan R, van den Akker J, et al. LEAP: Using machine learning to support variant classification in a clinical setting. Hum Mutat. 2020;41(6):1079–90.

312. Lai J, Yang J, Gamsiz Uzun ED, Rubenstein BM, Sarkar IN. LYRUS: a machine learning model for predicting the pathogenicity of missense variants. Bioinform Adv. 2022;2(1):vbab045.

313. Landrum MJ, Chitipiralla S, Brown GR, Chen C, Gu B, Hart J, et al. ClinVar: improvements to accessing data. Nucleic Acids Res. 2020;48(D1):D835–D44.

314. Landrum MJ, Kattman BL. ClinVar at five years: Delivering on the promise. Hum Mutat. 2018;39(11):1623–30.

315. Landrum MJ, Lee JM, Benson M, Brown GR, Chao C, Chitipiralla S, et al. ClinVar: improving access to variant interpretations and supporting evidence. Nucleic Acids Res. 2018;46(D1):D1062–D7.

316. Lek M, Karczewski KJ, Minikel EV, Samocha KE, Banks E, Fennell T, et al. Analysis of protein-coding genetic variation in 60,706 humans. Nature. 2016;536(7616):285-91.

317. Leman R, Gaildrat P, Le Gac G, Ka C, Fichou Y, Audrezet MP, et al. Novel diagnostic tool for prediction of variant spliceogenicity derived from a set of 395 combined in silico/in vitro studies: an international collaborative effort. Nucleic Acids Res. 2018;46(15):7913–23.

318. Leman R, Harter V, Atkinson A, Davy G, Rousselin A, Muller E, et al. SpliceLauncher: a tool for detection, annotation and relative quantification of alternative junctions from RNAseq data. Bioinformatics. 2020;36(5):1634–6.

319. Leman R, Parfait B, Vidaud D, Girodon E, Pacot L, Le Gac G, et al. SPiP: Splicing Prediction Pipeline, a machine learning tool for massive detection of exonic and intronic variant effects on mRNA splicing. Hum Mutat. 2022;43(12):2308–23.

320. Leslie R, O’Donnell CJ, Johnson AD. GRASP: analysis of genotype-phenotype results from 1390 genome-wide association studies and corresponding open access database. Bioinformatics. 2014;30(12):i185–94.

321. Li C, Zhi D, Wang K, Liu X. MetaRNN: differentiating rare pathogenic and rare benign missense SNVs and InDels using deep learning. Genome Med. 2022;14(1):115.

322. Li G, Pahari S, Murthy AK, Liang S, Fragoza R, Yu H, et al. SAAMBE-SEQ: a sequence-based method for predicting mutation effect on protein-protein binding affinity. Bioinformatics. 2021;37(7):992–9.

323. Li G, Panday SK, Alexov E. SAAFEC-SEQ: A Sequence-Based Method for Predicting the Effect of Single Point Mutations on Protein Thermodynamic Stability. Int J Mol Sci. 2021;22(2).

324. Li G, Yao S, Fan L. ProSTAGE: Predicting Effects of Mutations on Protein Stability by Using Protein Embeddings and Graph Convolutional Networks. J Chem Inf Model. 2024;64(2):340–7.

325. Li H, Liu S, Wang S, Zeng Q, Chen Y, Fang T, et al. Cancer SIGVAR: A semiautomated interpretation tool for germline variants of hereditary cancer-related genes. Hum Mutat. 2021;42(4):359–72.

326. Li M, Simonetti FL, Goncearenco A, Panchenko AR. MutaBind estimates and interprets the effects of sequence variants on protein-protein interactions. Nucleic Acids Res. 2016;44(W1):W494–501.

327. Liu Y, Dougherty JD. utr.annotation: a tool for annotating genomic variants that could influence post-transcriptional regulation. Bioinformatics. 2021;37(21):3926–8.

328. Lott MT, Leipzig JN, Derbeneva O, Xie HM, Chalkia D, Sarmady M, et al. mtDNA Variation and Analysis Using Mitomap and Mitomaster. Curr Protoc Bioinformatics. 2013;44(123):1 23 1-6.

329. Lou S, Cotter KA, Li T, Liang J, Mohsen H, Liu J, et al. GRAM: A GeneRAlized Model to predict the molecular effect of a non-coding variant in a cell-type specific manner. PLoS Genet. 2019;15(8):e1007860.

330. Lu H, Ma L, Quan C, Li L, Lu Y, Zhou G, et al. RegVar: Tissue-specific Prioritization of Non-coding Regulatory Variants. Genomics Proteomics Bioinformatics. 2023;21(2):385–95.

331. Malhis N, Jacobson M, Jones SJM, Gsponer J. LIST-S2: taxonomy based sorting of deleterious missense mutations across species. Nucleic Acids Res. 2020;48(W1):W154–W61.

332. Malhis N, Jones SJM, Gsponer J. Improved measures for evolutionary conservation that exploit taxonomy distances. Nat Commun. 2019;10(1):1556.

333. Markham JF, Yerneni S, Ryland GL, Leong HS, Fellowes A, Thompson ER, et al. CNspector: a web-based tool for visualisation and clinical diagnosis of copy number variation from next generation sequencing. Sci Rep. 2019;9(1):6426.

334. Marquet C, Heinzinger M, Olenyi T, Dallago C, Erckert K, Bernhofer M, et al. Embeddings from protein language models predict conservation and variant effects. Hum Genet. 2022;141(10):1629–47.

335. Martin-Antoniano I, Alonso L, Madrid M, Lopez de Maturana E, Malats N. DoriTool: A Bioinformatics Integrative Tool for Post-Association Functional Annotation. Public Health Genomics. 2017;20(2):126–35.

336. McVicker G, Gordon D, Davis C, Green P. Widespread genomic signatures of natural selection in hominid evolution. PLoS Genet. 2009;5(5):e1000471.

337. Menon R, Patel NV, Mohapatra A, Joshi CG. VDAP-GUI: a user-friendly pipeline for variant discovery and annotation of raw next-generation sequencing data. 3 Biotech. 2016;6(1):68.

338. Montanucci L, Capriotti E, Frank Y, Ben-Tal N, Fariselli P. DDGun: an untrained method for the prediction of protein stability changes upon single and multiple point variations. BMC Bioinformatics. 2019;20(Suppl 14):335.

339. Munro D, Singh M. DeMaSk: a deep mutational scanning substitution matrix and its use for variant impact prediction. Bioinformatics. 2021;36(22-23):5322–9.

340. Nachtegael C, Gravel B, Dillen A, Smits G, Nowe A, Papadimitriou S, et al. Scaling up oligogenic diseases research with OLIDA: the Oligogenic Diseases Database. Database (Oxford). 2022;2022.

341. Ng PC, Henikoff S. Predicting deleterious amino acid substitutions. Genome Res. 2001;11(5):863–74.

342. Nishio SY, Usami SI. The Clinical Next-Generation Sequencing Database: A Tool for the Unified Management of Clinical Information and Genetic Variants to Accelerate Variant Pathogenicity Classification. Hum Mutat. 2017;38(3):252–9.

343. Pagel KA, Antaki D, Lian A, Mort M, Cooper DN, Sebat J, et al. Pathogenicity and functional impact of non-frameshifting insertion/deletion variation in the human genome. PLoS Comput Biol. 2019;15(6):e1007112.

344. Pahari S, Li G, Murthy AK, Liang S, Fragoza R, Yu H, et al. SAAMBE-3D: Predicting Effect of Mutations on Protein-Protein Interactions. Int J Mol Sci. 2020;21(7).

345. Pais LS, Snow H, Weisburd B, Zhang S, Baxter SM, DiTroia S, et al. seqr: A web-based analysis and collaboration tool for rare disease genomics. Hum Mutat. 2022;43(6):698–707.

346. Palheta HGA, Goncalves WG, Brito LM, Dos Santos AR, Dos Reis Matsumoto M, Ribeiro-Dos-Santos A, et al. AmazonForest: In Silico Metaprediction of Pathogenic Variants. Biology (Basel). 2022;11(4).

347. Pancotti C, Benevenuta S, Repetto V, Birolo G, Capriotti E, Sanavia T, et al. A Deep-Learning Sequence-Based Method to Predict Protein Stability Changes Upon Genetic Variations. Genes (Basel). 2021;12(6).

348. Pei G, Hu R, Jia P, Zhao Z. DeepFun: a deep learning sequence-based model to decipher non-coding variant effect in a tissue-and cell type-specific manner. Nucleic Acids Res. 2021;49(W1):W131–W9.

349. Petukh M, Dai L, Alexov E. SAAMBE: Webserver to Predict the Charge of Binding Free Energy Caused by Amino Acids Mutations. Int J Mol Sci. 2016;17(4):547.

350. Pires DE, Ascher DB, Blundell TL. mCSM: predicting the effects of mutations in proteins using graph-based signatures. Bioinformatics. 2014;30(3):335–42.

351. Piriyapongsa J, Sukritha C, Kaewprommal P, Intarat C, Triparn K, Phornsiricharoenphant K, et al. PharmVIP: A Web-Based Tool for Pharmacogenomic Variant Analysis and Interpretation. J Pers Med. 2021;11(11).

352. Ponzoni L, Penaherrera DA, Oltvai ZN, Bahar I. Rhapsody: predicting the pathogenicity of human missense variants. Bioinformatics. 2020;36(10):3084–92.

353. Popov P, Bizin I, Gromiha M, A K, Frishman D. Prediction of disease-associated mutations in the transmembrane regions of proteins with known 3D structure. PLoS One. 2019;14(7):e0219452.

354. Prive F, Albinana C, Arbel J, Pasaniuc B, Vilhjalmsson BJ. Inferring disease architecture and predictive ability with LDpred2-auto. Am J Hum Genet. 2023;110(12):2042–55.

355. Prunier J, Lemacon A, Bastien A, Jafarikia M, Porth I, Robert C, et al. LD-annot: A Bioinformatics Tool to Automatically Provide Candidate SNPs With Annotations for Genetically Linked Genes. Front Genet. 2019;10:1192.

356. Qi H, Zhang H, Zhao Y, Chen C, Long JJ, Chung WK, et al. MVP predicts the pathogenicity of missense variants by deep learning. Nat Commun. 2021;12(1):510.

357. Quan L, Lv Q, Zhang Y. STRUM: structure-based prediction of protein stability changes upon single-point mutation. Bioinformatics. 2016;32(19):2936–46.

358. Quinodoz M, Peter VG, Bedoni N, Royer Bertrand B, Cisarova K, Salmaninejad A, et al. AutoMap is a high performance homozygosity mapping tool using next-generation sequencing data. Nat Commun. 2021;12(1):518.

359. Quinones-Valdez G, Fu T, Chan TW, Xiao X. scAllele: A versatile tool for the detection and analysis of variants in scRNA-seq. Sci Adv. 2022;8(35):eabn6398.

360. Radusky L, Modenutti C, Delgado J, Bustamante JP, Vishnopolska S, Kiel C, et al. VarQ: A Tool for the Structural and Functional Analysis of Human Protein Variants. Front Genet. 2018;9:620.

361. Raimondi D, Gazzo AM, Rooman M, Lenaerts T, Vranken WF. Multilevel biological characterization of exomic variants at the protein level significantly improves the identification of their deleterious effects. Bioinformatics. 2016;32(12):1797–804.

362. Raimondi D, Tanyalcin I, Ferte J, Gazzo A, Orlando G, Lenaerts T, et al. DEOGEN2: prediction and interactive visualization of single amino acid variant deleteriousness in human proteins. Nucleic Acids Res. 2017;45(W1):W201–W6.

363. Rastogi R, Stenson PD, Cooper DN, Bejerano G. X-CAP improves pathogenicity prediction of stopgain variants. Genome Med. 2022;14(1):81.

364. Rathinakannan VS, Schukov HP, Heron S, Schleutker J, Sipeky C. ShAn: An easy-to-use tool for interactive and integrated variant annotation. PLoS One. 2020;15(7):e0235669.

365. Ravichandran V, Shameer Z, Kemel Y, Walsh M, Cadoo K, Lipkin S, et al. Toward automation of germline variant curation in clinical cancer genetics. Genet Med. 2019;21(9):2116–25.

366. Rehm HL, Berg JS, Brooks LD, Bustamante CD, Evans JP, Landrum MJ, et al. ClinGen--the Clinical Genome Resource. N Engl J Med. 2015;372(23):2235–42.

367. Rentzsch P, Witten D, Cooper GM, Shendure J, Kircher M. CADD: predicting the deleteriousness of variants throughout the human genome. Nucleic Acids Res. 2019;47(D1):D886–D94.

368. Rives A, Meier J, Sercu T, Goyal S, Lin Z, Liu J, et al. Biological structure and function emerge from scaling unsupervised learning to 250 million protein sequences. Proc Natl Acad Sci U S A. 2021;118(15).

369. Rodrigues CH, Pires DE, Ascher DB. DynaMut: predicting the impact of mutations on protein conformation, flexibility and stability. Nucleic Acids Res. 2018;46(W1):W350–W5.

370. Rodrigues CHM, Pires DEV, Ascher DB. DynaMut2: Assessing changes in stability and flexibility upon single and multiple point missense mutations. Protein Sci. 2021;30(1):60–9.

371. Rogers MF, Shihab HA, Gaunt TR, Campbell C. CScape: a tool for predicting oncogenic single-point mutations in the cancer genome. Sci Rep. 2017;7(1):11597.

372. Rogers MF, Shihab HA, Mort M, Cooper DN, Gaunt TR, Campbell C. FATHMM-XF: accurate prediction of pathogenic point mutations via extended features. Bioinformatics. 2018;34(3):511–3.

373. Sasorith S, Baux D, Bergougnoux A, Paulet D, Lahure A, Bareil C, et al. The CYSMA web server: An example of integrative tool for in silico analysis of missense variants identified in Mendelian disorders. Hum Mutat. 2020;41(2):375–86.

374. Seva J, Wiegandt DL, Gotze J, Lamping M, Rieke D, Schafer R, et al. VIST-a Variant-Information Search Tool for precision oncology. BMC Bioinformatics. 2019;20(1):429.

375. Shamsi Z, Chan M, Shukla D. TLmutation: Predicting the Effects of Mutations Using Transfer Learning. J Phys Chem B. 2020;124(19):3845–54.

376. Sharo AG, Hu Z, Sunyaev SR, Brenner SE. StrVCTVRE: A supervised learning method to predict the pathogenicity of human genome structural variants. Am J Hum Genet. 2022;109(2):195–209.

377. Shibata A, Okuno T, Rahman MA, Azuma Y, Takeda J, Masuda A, et al. IntSplice: prediction of the splicing consequences of intronic single-nucleotide variations in the human genome. J Hum Genet. 2016;61(7):633–40.

378. Shin J, Jeon J, Jung D, Kim K, Kim YJ, Jeong DH, et al. PhenGenVar: A User-Friendly Genetic Variant Detection and Visualization Tool for Precision Medicine. J Pers Med. 2022;12(6).

379. Sokolova K, Theesfeld CL, Wong AK, Zhang Z, Dolinski K, Troyanskaya OG. Atlas of primary cell-type-specific sequence models of gene expression and variant effects. Cell Rep Methods. 2023;3(9):100580.

380. Spector JD, Wiita AP. ClinTAD: a tool for copy number variant interpretation in the context of topologically associated domains. J Hum Genet. 2019;64(5):437–43.

381. Staley JR, Blackshaw J, Kamat MA, Ellis S, Surendran P, Sun BB, et al. PhenoScanner: a database of human genotype-phenotype associations. Bioinformatics. 2016;32(20):3207–9.

382. Steinhaus R, Proft S, Schuelke M, Cooper DN, Schwarz JM, Seelow D. MutationTaster2021. Nucleic Acids Res. 2021;49(W1):W446-W51.

383. Stenson PD, Ball EV, Mort M, Phillips AD, Shaw K, Cooper DN. The Human Gene Mutation Database (HGMD) and its exploitation in the fields of personalized genomics and molecular evolution. Curr Protoc Bioinformatics. 2012;Chapter 1:1 13 1-1 20.

384. Stenson PD, Ball EV, Mort M, Phillips AD, Shiel JA, Thomas NS, et al. Human Gene Mutation Database (HGMD): 2003 update. Hum Mutat. 2003;21(6):577–81.

385. Stenson PD, Mort M, Ball EV, Chapman M, Evans K, Azevedo L, et al. The Human Gene Mutation Database (HGMD((R))): optimizing its use in a clinical diagnostic or research setting. Hum Genet. 2020;139(10):1197–207.

386. Stenson PD, Mort M, Ball EV, Howells K, Phillips AD, Thomas NS, et al. The Human Gene Mutation Database: 2008 update. Genome Med. 2009;1(1):13.

387. Stenson PD, Mort M, Ball EV, Shaw K, Phillips A, Cooper DN. The Human Gene Mutation Database: building a comprehensive mutation repository for clinical and molecular genetics, diagnostic testing and personalized genomic medicine. Hum Genet. 2014;133(1):1–9.

388. Sun W, Duan T, Ye P, Chen K, Zhang G, Lai M, et al. TSVdb: a web-tool for TCGA splicing variants analysis. BMC Genomics. 2018;19(1):405.

389. Sundaram L, Gao H, Padigepati SR, McRae JF, Li Y, Kosmicki JA, et al. Predicting the clinical impact of human mutation with deep neural networks. Nat Genet. 2018;50(8):1161–70.

390. Takata A, Hamanaka K, Matsumoto N. Refinement of the clinical variant interpretation framework by statistical evidence and machine learning. Med. 2021;2(5):611-32 e9.

391. Takeda JI, Fukami S, Tamura A, Shibata A, Ohno K. IntSplice2: Prediction of the Splicing Effects of Intronic Single-Nucleotide Variants Using LightGBM Modeling. Front Genet. 2021;12:701076.

392. Takeda JI, Nanatsue K, Yamagishi R, Ito M, Haga N, Hirata H, et al. InMeRF: prediction of pathogenicity of missense variants by individual modeling for each amino acid substitution. NAR Genom Bioinform. 2020;2(2):lqaa038.

393. Tamborero D, Rubio-Perez C, Deu-Pons J, Schroeder MP, Vivancos A, Rovira A, et al. Cancer Genome Interpreter annotates the biological and clinical relevance of tumor alterations. Genome Med. 2018;10(1):25.

394. Thanapattheerakul T, Engchuan W, Chan JH. Predicting the effect of variants on splicing using Convolutional Neural Networks. PeerJ. 2020;8:e9470.

395. Thornton AM, Fang L, Lo A, McSharry M, Haan D, O’Brien C, et al. eVIP2: Expression-based variant impact phenotyping to predict the function of gene variants. PLoS Comput Biol. 2021;17(7):e1009132.

396. Tokheim C, Karchin R. CHASMplus Reveals the Scope of Somatic Missense Mutations Driving Human Cancers. Cell Syst. 2019;9(1):9–23 e8.

397. Tong SY, Fan K, Zhou ZW, Liu LY, Zhang SQ, Fu Y, et al. mvPPT: A Highly Efficient and Sensitive Pathogenicity Prediction Tool for Missense Variants. Genomics Proteomics Bioinformatics. 2023;21(2):414–26.

398. Trovato A, Seno F, Tosatto SC. The PASTA server for protein aggregation prediction. Protein Eng Des Sel. 2007;20(10):521–3.

399. Turner TN, Yi Q, Krumm N, Huddleston J, Hoekzema K, HA FS, et al. denovo-db: a compendium of human de novo variants. Nucleic Acids Res. 2017;45(D1):D804–D11.

400. Wang J, Liu Z, Bellen HJ, Yamamoto S. Navigating MARRVEL, a Web-Based Tool that Integrates Human Genomics and Model Organism Genetics Information. J Vis Exp. 2019(150).

401. Wang M, Deng W, Samuels DC, Zhao Z, Simon LM. MitoTrace: A Computational Framework for Analyzing Mitochondrial Variation in Single-Cell RNA Sequencing Data. Genes (Basel). 2023;14(6).

402. Ward LD, Kellis M. HaploReg: a resource for exploring chromatin states, conservation, and regulatory motif alterations within sets of genetically linked variants. Nucleic Acids Res. 2012;40(Database issue):D930-4.

403. Wells A, Heckerman D, Torkamani A, Yin L, Sebat J, Ren B, et al. Ranking of non-coding pathogenic variants and putative essential regions of the human genome. Nat Commun. 2019;10(1):5241.

404. Won DG, Kim DW, Woo J, Lee K. 3Cnet: pathogenicity prediction of human variants using multitask learning with evolutionary constraints. Bioinformatics. 2021;37(24):4626–34.

405. Woodard J, Zhang C, Zhang Y. ADDRESS: A Database of Disease-associated Human Variants Incorporating Protein Structure and Folding Stabilities. J Mol Biol. 2021;433(11):166840.

406. Wu Y, Li R, Sun S, Weile J, Roth FP. Improved pathogenicity prediction for rare human missense variants. Am J Hum Genet. 2021;108(10):1891–906.

407. Xavier A, Scott RJ, Talseth-Palmer BA. TAPES: A tool for assessment and prioritisation in exome studies. PLoS Comput Biol. 2019;15(10):e1007453.

408. Xiang J, Peng J, Baxter S, Peng Z. AutoPVS1: An automatic classification tool for PVS1 interpretation of null variants. Hum Mutat. 2020;41(9):1488–98.

409. Xiao Y, Wang J, Li J, Zhang P, Li J, Zhou Y, et al. An analytical framework for decoding cell type-specific genetic variation of gene regulation. Nat Commun. 2023;14(1):3884.

410. Yue Z, Zhao L, Cheng N, Yan H, Xia J. dbCID: a manually curated resource for exploring the driver indels in human cancer. Brief Bioinform. 2019;20(5):1925–33.

411. Zhang H, Xu MS, Fan X, Chung WK, Shen Y. Predicting functional effect of missense variants using graph attention neural networks. Nat Mach Intell. 2022;4(11):1017–28.

412. Zhang N, Chen Y, Lu H, Zhao F, Alvarez RV, Goncearenco A, et al. MutaBind2: Predicting the Impacts of Single and Multiple Mutations on Protein-Protein Interactions. iScience. 2020;23(3):100939.

413. Zhang X, Walsh R, Whiffin N, Buchan R, Midwinter W, Wilk A, et al. Disease-specific variant pathogenicity prediction significantly improves variant interpretation in inherited cardiac conditions. Genet Med. 2021;23(1):69–79.

414. Zhou J, Gao J, Zhang H, Zhao D, Li A, Iqbal F, et al. PedMiner: a tool for linkage analysis-based identification of disease-associated variants using family based whole-exome sequencing data. Brief Bioinform. 2021;22(3).

415. Zia M, Spurgeon P, Levesque A, Furlani T, Wang J. GenESysV: a fast, intuitive and scalable genome exploration open source tool for variants generated from high-throughput sequencing projects. BMC Bioinformatics. 2019;20(1):61.

416. Rastogi R, Chung R, Li S, Li C, Lee K, Woo J, et al. Critical assessment of missense variant effect predictors on disease-relevant variant data. bioRxiv. 2024.

